# Genetic requirements for cell division in a genomically minimal cell

**DOI:** 10.1101/2020.10.07.326892

**Authors:** James F. Pelletier, Lijie Sun, Kim S. Wise, Nacyra Assad-Garcia, Bogumil J. Karas, Thomas J. Deerinck, Mark H. Ellisman, Andreas Mershin, Neil Gershenfeld, Ray-Yuan Chuang, John I. Glass, Elizabeth A. Strychalski

**Affiliations:** Center for Bits and Atoms, Massachusetts Institute of Technology, Cambridge, MA 02139, USA; National Institute of Standards and Technology, Gaithersburg, MD 20899, USA; J. Craig Venter Institute, La Jolla, CA 92037, USA; Department of Biochemistry, Schulich School of Medicine and Dentistry, The University of Western Ontario, London, ON, N6A 5C1, Canada; National Center for Microscopy and Imaging Research, University of California–San Diego, La Jolla, CA 92037, USA

## Abstract

Genomically minimal cells, such as JCVI-syn3.0, offer a platform to clarify genes underlying core physiological processes. While this minimal cell includes genes essential for population growth, the physiology of its single cells remained uncharacterized. To investigate striking morphological variation in JCVI-syn3.0 cells, we present an approach to characterize cell propagation and determine genes affecting cell morphology. Microfluidic chemostats allowed observation of intrinsic cell dynamics resulting in irregular morphologies. The addition of 19 genes not retained in JCVI-syn3.0 generated JCVI-syn3A, which presents significantly less morphological variation than JCVI-syn3.0. We further identified seven of these 19 genes, including two known cell division genes *ftsZ* and *sepF* and five genes of unknown function, required together to restore cell morphology and division similar to JCVI-syn1.0. This surprising result emphasizes the polygenic nature of cell morphology, as well as the importance of a Z-ring and membrane properties in the physiology of genomically minimal cells.

## Introduction

Synthetic and engineering biology are creating new capabilities to investigate and program fundamental biological processes, for example, from sensors programmed as genetic circuits enabling control (Chen et al., 2020; Nielsen et al., 2016), to organisms with wholly recoded genomes (Annaluru et al., 2014; Fredens et al., 2019; Ostrov et al., 2016), to synthetic cells constructed from non-living parts (Hürtgen et al., 2019; Noireaux and Liu, 2020). Genome minimization offers a compelling synthetic biology approach to study the emergence of fundamental physiological processes from interactions between essential genes. Towards this goal, researchers at the J. Craig Venter Institute (JCVI) and collaborators applied genome minimization and an engineering biology workflow to develop a tractable platform for unicellular life that reflects known organisms and comprises the simplest free-living system. They accomplished this, by building a functional synthetic genome that drives the propagation of a free-living cell (JCVI-syn1.0, nearly wild type *Mycoplasma mycoides* subspecies *capri*) (Gibson et al., 2010) and subsequently reducing genome complexity to deliver a nearly minimal living bacterium, JCVI-syn3.0 (Figs 1A-C) (Hutchison et al., 2016). Genome minimization leveraged an engineering design-build-test-learn workflow, based on the evaluation and combinatorial assembly of modular genomic segments, as well as empirical results obtained from transposon mutagenesis to inform gene deletions. This approach both reduced bias in identifying essential genes and provided tools to alter, rebuild, and investigate the genome and encoded functions. With only 473 genes, the resulting strain JCVI-syn3.0 boasts the smallest genome of any free-living organism; yet, 149 of these essential genes are classified as genes of unknown or generic function. Many of these genes are conserved in walled bacteria, from which mycoplasmas evolved through massive gene loss. Mycoplasmas, such as JCVI-syn1.0 and JCVI-syn3.0, thus offer enabling platforms to probe essential processes conserved broadly in cellular life, as demonstrated here.

**Figure 1.**
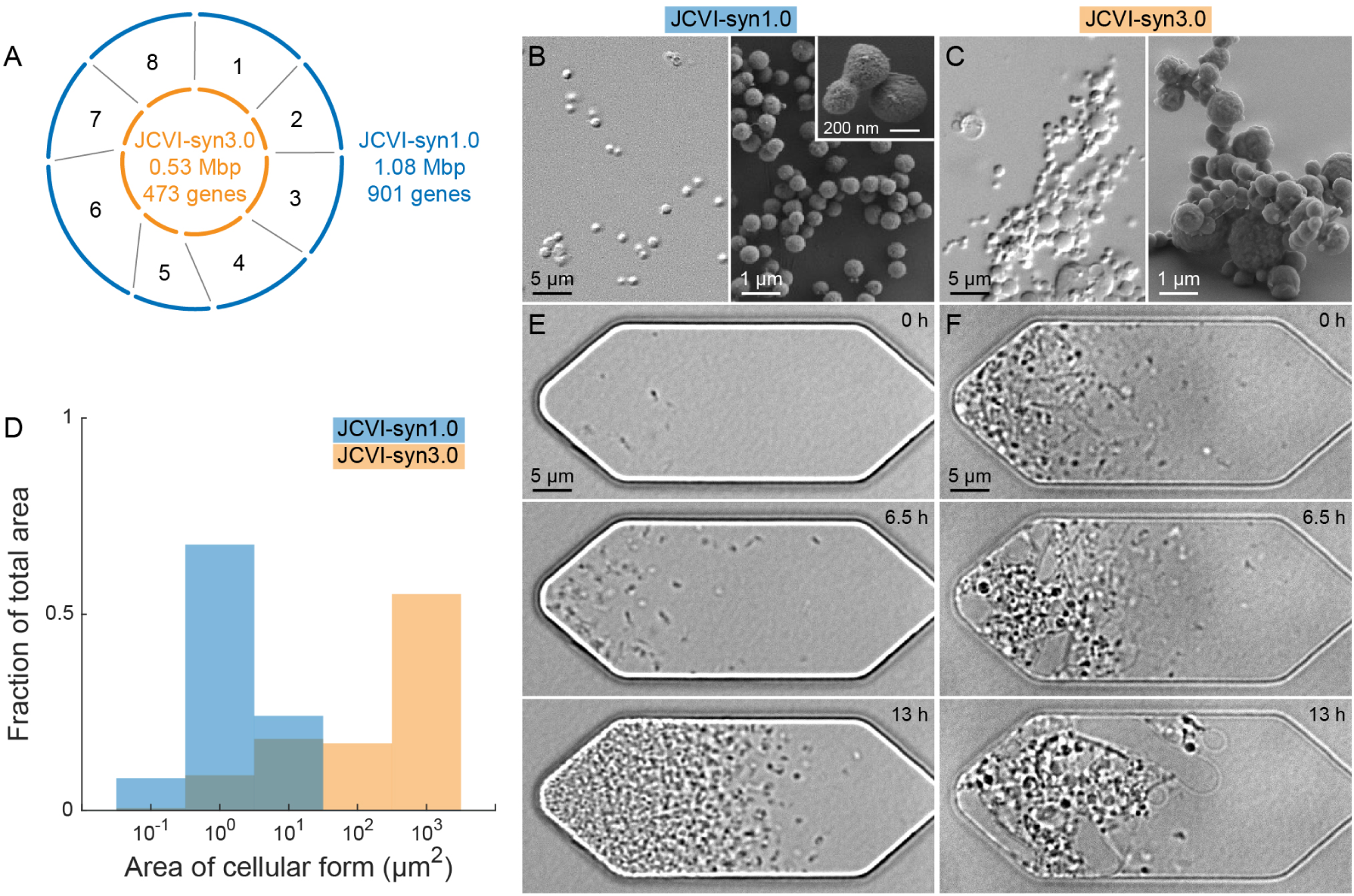
Genome reduction resulted in morphological variation at the cellular scale. (A) JCVI-syn3.0 is a genomically minimized cell derived from JCVI-syn1.0, which is similar to wild type *M. mycoides*. Each genomic segment was minimized independently and then reassembled to produce the minimal JCVI-syn3.0 genome. (B, C) Representative optical and scanning electron micrographs of cells grown in bulk liquid culture show (B) JCVI-syn1.0 with submicron, round cells of uniform size and shape, and (C) JCVI-syn3.0 with variable size and shape. Some JCVI-syn3.0 cells appeared similar to JCVI-syn1.0 cells, while others clustered together or included large cellular forms with diameters >10 µm or irregular shapes. (D) Frequency histograms of the area of each cell or cellular form show clear differences in the overall morphological variation of these strains at the cellular scale. (E, F) Timelapse optical micrographs of culture in microfluidic chemostats revealed the dynamic propagation of normal JCVI-syn1.0 cells and pleomorphic JCVI-syn3.0 cellular forms.

JCVI-syn3.0 grows axenically in a complex liquid medium that provides many nutrients the cell cannot synthesize, similar to other mycoplasmas. A minimal replicative unit passes through a 0.22-micron filter and forms a typical mycoplasma colony. Moreover, measurements of cell-associated nucleic acid indicate logarithmic growth (Hutchison et al., 2016). The cell, then, meets two *sine qua non* of life: propagation of a membrane-bound compartment containing the genome with its replicative machinery and replication of that genome. Nevertheless, a wholly unexpected feature of JCVI-syn3.0 is the striking morphological variation of individual cells, with filamentous, vesicular, and other irregular forms described previously using static optical and scanning electron micrographs (Fig 1C) (Hutchison et al., 2016). These morphologies are absent in both wild type *M. mycoides* and JCVI-syn1.0 cells, which exhibit what we refer to here as “normal morphology” consisting of spherical cells ≈400 nm in diameter (Fig 1B) (Hutchison et al., 2016).

Static images cannot reveal the biogenesis of pleomorphic cellular forms, their content, or their relevance in propagation. We report here on the propagation of membrane compartments and confirm these shapes arise from morphological dynamics intrinsic to individual cells via imaging in novel microfluidic chemostats. We then investigated the genetic basis of the pleomorphic phenotype using a reverse genetics approach to restore normal morphology through various, nearly minimal genomes. The resulting nearly minimal strain, JCVI-syn3A, has 19 additional genes and presents significantly less morphological variation than JCVI-syn3.0. JCVI-syn3A is mechanically robust to the liquid handling required for biological research and compatible with practical computational modeling of a minimal metabolism (Breuer et al., 2019). Here, we determined that seven of the 19 genes, of known and unknown function, were necessary to restore normal morphology. The surprising requirement for all seven genes, including *ftsZ, sepF*, and others of unknown function, highlights the importance of both a Z-ring and membrane biophysics in the scission of genomically minimized cells.

## Results

### Dynamic propagation of JCVI-syn3.0

To characterize the morphology of JCVI-syn3.0 and related genomically minimized strains at the cellular scale, we obtained timelapse images in microfluidic chemostats (Figs 1E,F, SI Movies 1,2). This platform shielded cells from shear flow, which can affect mycoplasmal morphology (Razin et al., 1967) and has been proposed to facilitate scission of primordial cells (Chen, 2009). Thus, the microfluidic chemostats isolated mechanisms of propagation and cell division intrinsic to a cell and revealed directly the emergence and dynamics of diverse morphologies (Methods, Fig S1). Notably, strains exhibited similar phenotypic variation in both static liquid culture and microfluidic chemostats.

Using these observations, we separated strains into two general morphological classes: normal and pleomorphic. JCVI-syn1.0 defined the normal morphology, with round, submicron cells, largely disconnected from one another (Fig 1B, SI Movie 1). In contrast, JCVI-syn3.0 exemplified pleomorphic morphologies, including large cellular forms with diameters ≥10 µm and irregular shapes (Fig 1C, SI Movie 2). We use the term “cellular form” to describe complex morphologies with adjacent substructures. While some JCVI-syn3.0 cells appeared similar to JCVI-syn1.0 cells, the majority consisted of large cellular forms (Fig 1C, SI Movie 2). The area distribution of cells and cellular forms was quantified using static liquid culture (Fig 1D, Methods), while the dynamics of propagation were observed in microfluidic chemostats (Figs 1E,F). We apply the general term “propagation” to describe shape changes that accompany growth and reserve the term “cell division” for complete scission.

### Morphological diversity from a single minimized genomic segment

To perform an unbiased screen for genes correlated with morphological variation, we exploited the segmented structure of the JCVI-syn3.0 genome. During the process of genome minimization from JCVI-syn1.0 to JCVI-syn3.0, genes or gene clusters were removed in various combinations from individual genomic segments. Each minimized segment in the context of an otherwise JCVI-syn1.0 genome showed viability and growth at the population scale (Hutchison et al., 2016). Re-examining these strains here at the cellular scale, we noted that one strain, termed RGD6 (<u>R</u>educed <u>G</u>enome <u>D</u>esign segment <u>6</u>), contained the fully minimized genomic segment 6 and demonstrated striking pleomorphism similar to JCVI-syn3.0 (Fig 2, cf. Fig 1C). Each of the other minimized segments in an otherwise JCVI-syn1.0 genome resulted in normal or nearly normal morphology (Fig 2).

**Figure 2.**
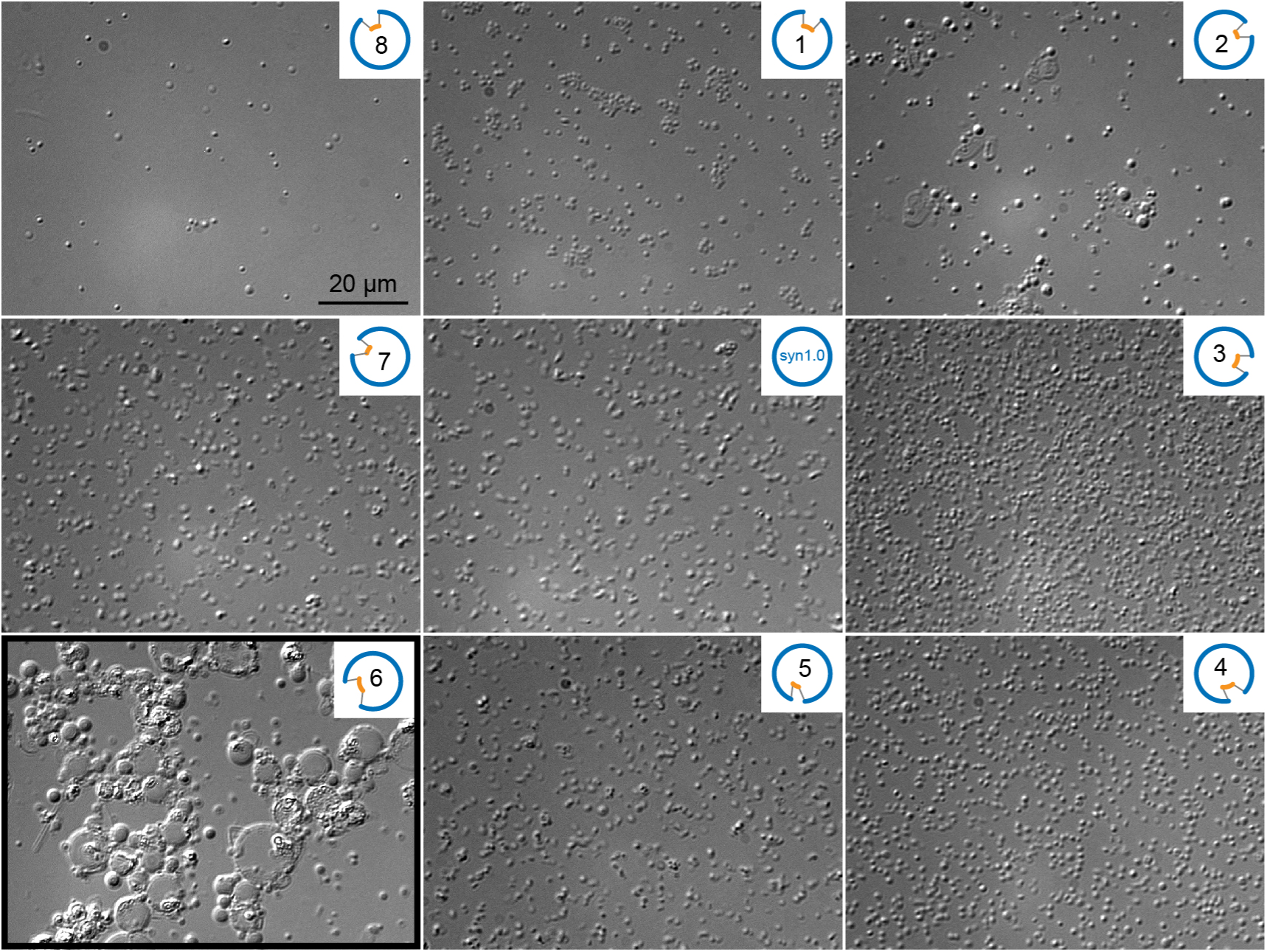
Genes in segment 6 strongly influenced morphological variation at the cellular scale. Tests of each minimized genomic segment in the context of a JCVI-syn1.0 genome revealed that segment 6 generated pleomorphic cellular forms similar to JCVI-syn3.0. Samples were prepared and imaged as for Figs 1B,C.

The comparable growth rate of RGD6 to JCVI-syn1.0 facilitated experimental investigation of the propagation of pleomorphic cellular forms (Fig 3, SI Movies 4,5). JCVI-syn3.0 appeared similar to RGD6, including irregular and filamentous cellular forms, which exhibited branching or pearling, with round, connected substructures (Fig 3B, SI Movie 3). In a representative microfluidic experiment, a single RGD6 propagated into a striking variety of morphologies (Fig 3A, SI Movie 4). The small cell grew into a filamentous cellular form, and vesicles appeared at the ends (Fig 3A). The vesicles lacked observable mCherry (a marker for cytoplasmic protein) from their first appearance. The filamentous cellular form increased in length and bent to fit within the chamber, while the vesicles continued to grow. Note that not all RGD6 cells showed filamentous propagation (cf. Figs 2,4, and SI Movie 5).

**Figure 3.**
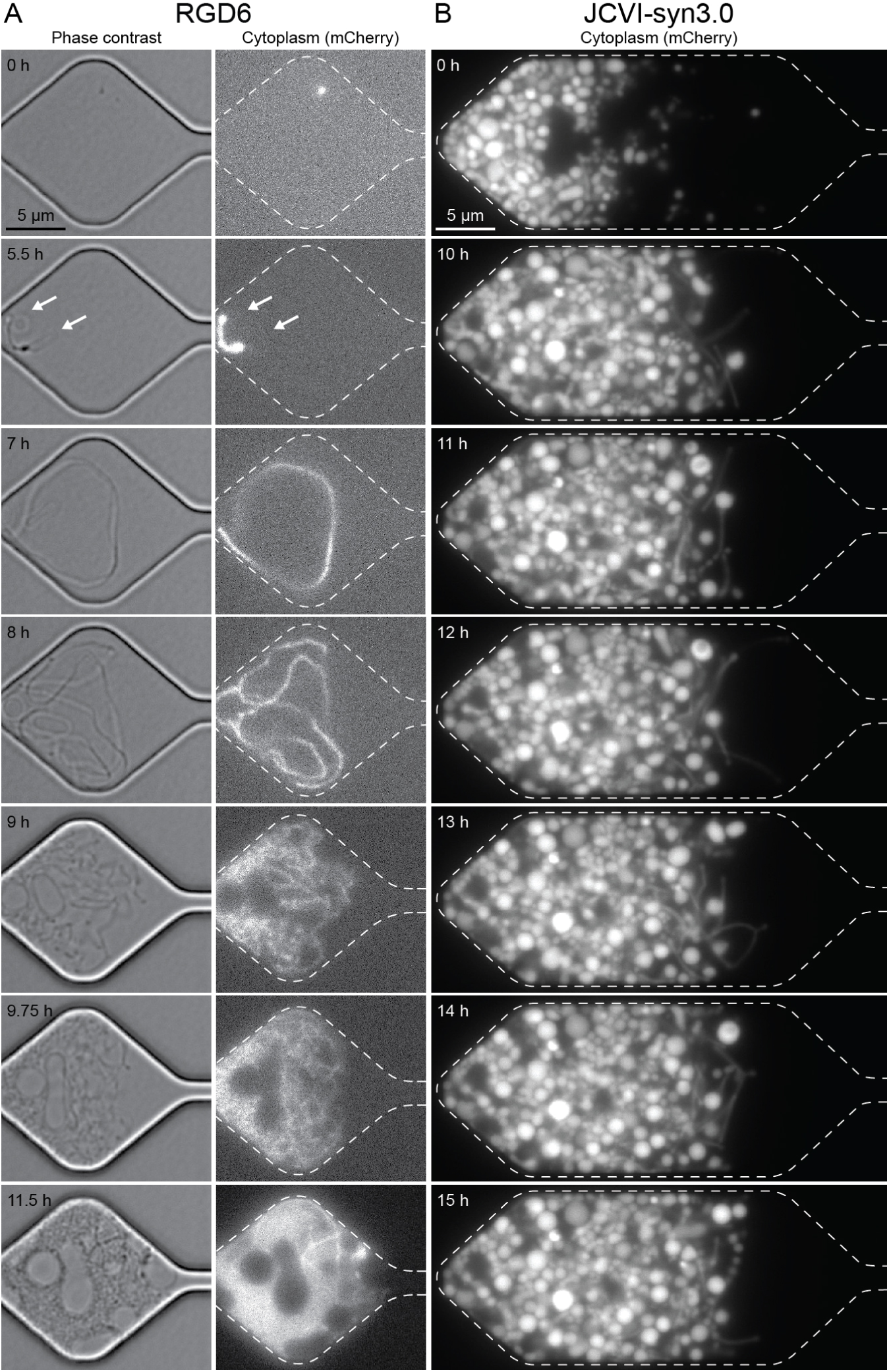
RGD6 and JCVI-syn3.0 exhibited filamentation, branching, pearling, and other morphological dynamics in the absence of shear flow. (A) RGD6 was capable of filamentous propagation in the shear-free environment of microfluidic chemostats. The left column shows phase contrast images, while the right column shows constitutively expressed mCherry as a marker for cytoplasm. Chemostat walls are indicated as dotted lines. White arrows at 5.5 h indicate the appearance of vesicles at the ends of the filament. Vesicles lacked mCherry from their first appearance. (B) Growth of JCVI-syn3.0 produces pearled and branched filaments, along with other morphologies, shown here in a representative chemostat originally loaded with many cells.

**Figure 4.**
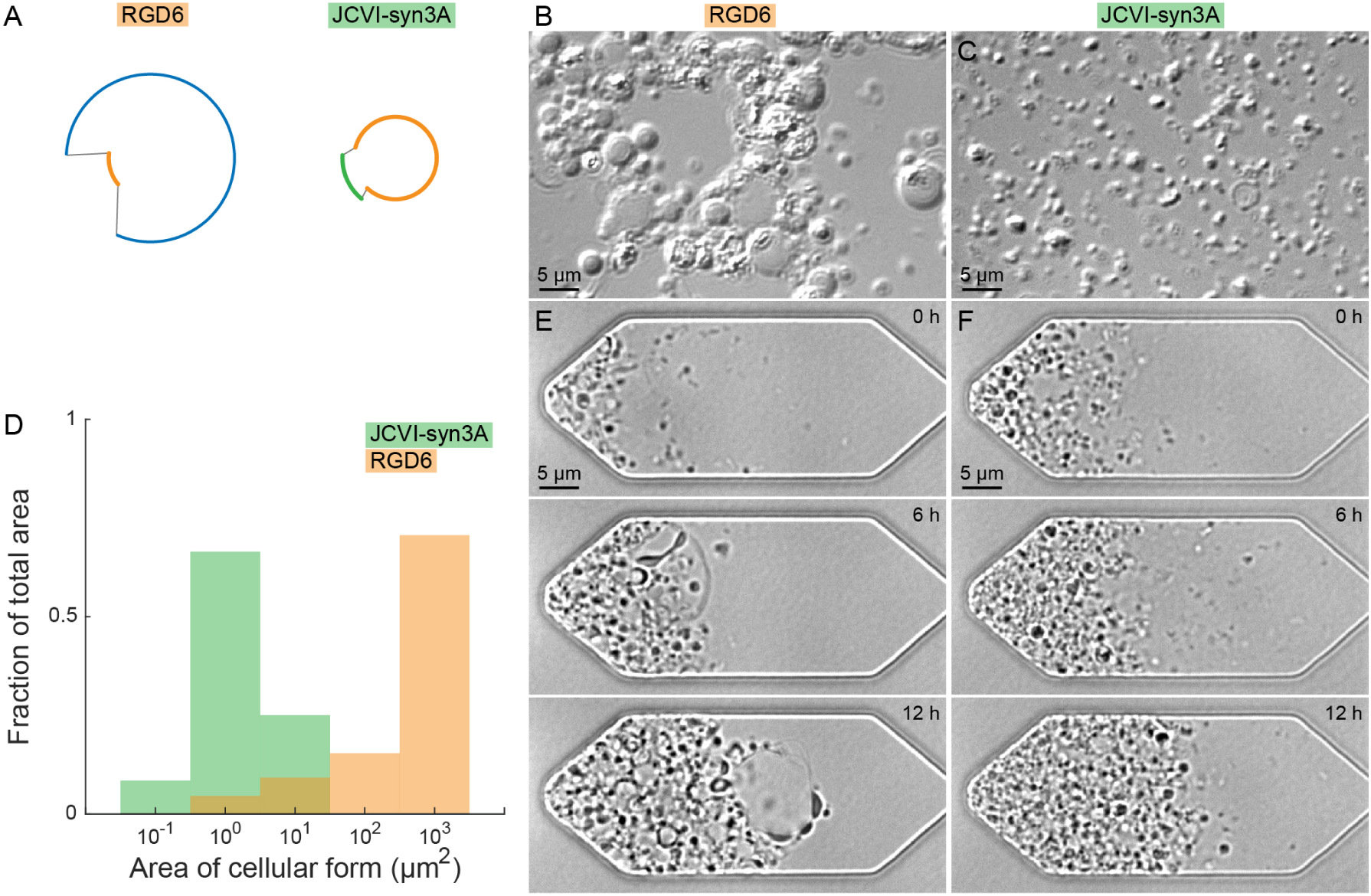
Addition of 19 genes to JCVI-syn3.0 significantly reduced morphological variation. (A) RGD6 includes the minimized segment 6 from JCVI-syn3.0, while JCVI-syn3A has a nearly minimal segment 6 containing 19 additional genes. (B, C) Optical micrographs show cells grown in bulk liquid culture. (B) RGD6 produced dramatic morphological variation similar to JCVI-syn3.0. (C) In contrast, JCVI-syn3A exhibited significantly less morphological variation, similar to JCVI-syn1.0. (D) The growth of JCVI-syn3A in the microfluidic chemostat appeared similar to that of JCVI-syn1.0 (Fig 1E). The significant reduction in morphological variation between RGD6 and JCVI-syn3A is evident by comparing (B,C) optical micrographs, (D) the frequency histograms of the area of cellular forms, and (E, F) growth in microfluidic chemostats.

To probe the internal structure of pleomorphic cellular forms, fluorescence imaging informed cellular composition and organization. We visualized cytoplasm with constitutively expressed fluorescent mCherry protein, nucleoids with Hoechst 33342 stain, membranes with the lipophilic dye SP-DiOC18(3), and extracellular growth medium with fluorophore-conjugated dextran (Fig S2). Filamentous cellular forms contained mCherry and often appeared with multiple nucleoids along their length, suggesting genome replication and segregation continued in the absence of complete scission. Although vesicles generally did not contain observable mCherry, they were labeled by the membrane dye and excluded fluorescent dextran from their interiors (Fig S2). This finding suggests that vesicles lacked cellular machinery needed for mCherry expression but remained bounded by membranes impermeable to macromolecules.

### Genomic restoration of normal morphology in a nearly minimal cell

To determine which genes in segment 6 are required for normal division, we leveraged genetics approaches enabled by the synthetic, modular, and minimal aspects of the genome. In particular, we pursued a systematic, reverse genetics approach, testing the dependence of morphological variation first on gene clusters and then on individual genes in segment 6. Among the 76 genes removed from segment 6 to generate JCVI-syn3.0 was an *ftsZ*-containing cluster (genes *520-522*), which lay within the *dcw* locus, a conserved cluster of genes known to participate in programmed cell division in most bacteria (Alarcón et al., 2007; Benders et al., 2005; Eraso et al., 2014; Vedyaykin et al., 2019). One nearly minimal version of segment 6 retained this *ftsZ*-containing cluster, along with genes outside the *dcw* cluster. This segment conferred a nearly normal morphology, when replacing the minimized segment 6 in JCVI-syn3.0, thereby creating a strain JCVI-syn3A (Fig 4A, SI Movie 6). The < 2 h doubling time of JCVI-syn3A is reduced from that of JCVI-syn3.0 (Breuer et al., 2019). The fully annotated genome sequence of JCVI-syn3A is deposited in NCBI (GenBank CP016816.2).

Segment 6 in JCVI-syn3A contains 19 genes not present in the fully minimized version of JCVI-syn3.0 (Fig 5). We proceeded to determine those genes required for normal morphology by adding subsets of these genes to JCVI-syn3.0. Three of these genes represent a redundant copy of the rRNA operon, which we determine here to not affect phenotype. The remaining 16 genes encoding for proteins were grouped in eight clusters of contiguous genes (Fig 5A). In principle, clusters could represent single transcriptional units. To account for this, we introduced clusters of contiguous genes into JCVI-syn3.0 and then followed this workflow by deleting individual genes. Genes or gene clusters were reintroduced at ectopic loci and included adjoining sequence to retain native regulatory regions (Hutchison et al., 2016). These precautions allowed us to evaluate whether the presence of a gene or cluster, even outside its native locus, was sufficient to confer the normal phenotype.

**Figure 5.**
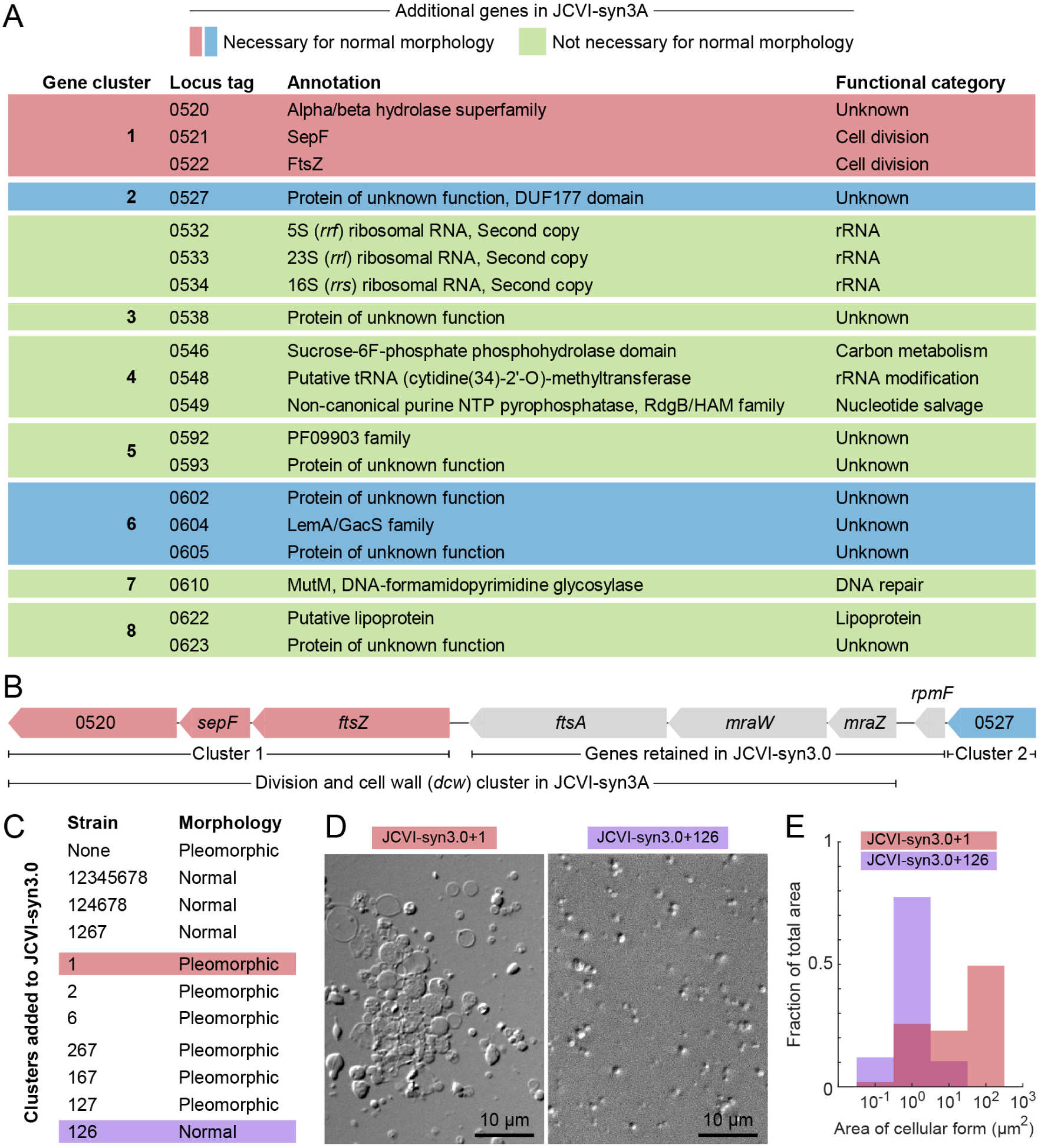
Seven of the 19 additional genes in JCVI-syn3A restored normal morphology to JCVI-syn3.0. (A) 19 additional genes in JCVI-syn3A are not retained in JCVI-syn3.0 and occurred in clusters of single or contiguous genes. One cluster represents a second copy of an rRNA operon. The other eight clusters encode proteins. (B) The *ftsZ*-containing cluster 1 is contained within a highly conserved division and cell wall (*dcw*) cluster in JCVI-syn3A, part of which is retained in JCVI-syn3.0. Arrows indicate gene direction, and black lines indicate intergenic sequences. (C) The addition of cluster 1 alone was insufficient to restore normal morphology, while clusters 1, 2, and 6 together, which include genes of unknown function, recovered the normal phenotype. (D) Optical micrographs show the pleomorphism of JCVI-syn3.0+1 and the normal morphology of JCVI-syn3.0+126. Optical micrographs of all strains are shown in Fig S4. (E) Area distributions of cellular forms confirm these morphological classifications.

We hypothesized that addition of the *ftsZ*-containing cluster would restore normal cell division and would alone confer the same reduction in morphological variation as observed in JCVI-syn3A. The *ftsZ*-containing cluster comprises three genes – *ftsZ, sepF*, and an adjacent hydrolase of unknown substrate specificity. Although part of the highly conserved division and cell wall (*dcw*) cluster present in the original JCVI-syn1.0 (Fig 5B), this subcluster was removed during genome minimization. Surprisingly, adding the *ftsZ*-containing cluster alone was not sufficient to reduce morphological variation in JCVI-syn3.0 (Figs 5C,D). Moreover, JCVI-syn3A retained a nearly normal morphology upon deletion of the *ftsZ*-containing cluster (Fig S4). The unexpectedly complex genetic basis for this phenotype required that we apply a more systematic approach to identify genes necessary and sufficient to restore the more normal phenotype in the JCVI-syn3.0 genomic context.

We validated our approach of adding genes to ectopic loci by adding all eight clusters to the JCVI-syn3.0 genome and found this restored normal morphology (Fig 5C, strain JCVI-syn3.0+12345678). This strain had the same gene content as JCVI-syn3A minus the redundant rRNA operon, thereby confirming that this second rRNA operon was not necessary for the normal phenotype. We then included increasingly fewer clusters, until we arrived at a construct containing only four of the clusters (strain JCVI-syn3.0+1267). Based on this strain, we concluded the four clusters omitted (3, 4, 5, and 8) were also not necessary for the normal morphology. We next added clusters individually to JCVI-syn3.0 and found that all strains remained pleomorphic, indicating each cluster alone was insufficient to restore the normal phenotype. We thus attempted omitting each cluster from JCVI-syn3.0+1267. While cluster 7 was dispensable, all others were necessary to restore the normal morphology.

Having determined all necessary clusters, we tested each gene individually. Employing the genome of strain JCVI-syn3.0+126 as a framework, we deleted each of the seven genes. These included the three genes in the *dcw* cluster, as well as the four genes of unknown function outside this locus. Gene coding regions were deleted in-frame to preserve transcriptional and translational integrity for genes within possible operons. None of the seven genes could be removed without reverting to the pleomorphic phenotype of JCVI-syn3.0 (Table S4, Fig S4). This provides strong evidence that all seven genes were necessary and together sufficient to reduce morphological variation in JCVI-syn3.0 to that of JCVI-syn3A (Figs 5D,E).

## Discussion

### Morphological dynamics intrinsic to cells revealed by microfluidic chemostats

As mycoplasmas lack a peptidoglycan cell wall, mycoplasmal morphologies are sensitive to shear flow, which can drive filamentation (Razin et al., 1967). Here, the microfluidic chemostat sheltered cells from shear flow, while enabling observation of their growth and propagation and ensuring that filamentous and other forms reflected morphological dynamics intrinsic to cells. In striking contrast to JCVI-syn1.0, our study suggests that JCVI-syn3.0 has lost the ability to control cell division. This is seen perhaps most clearly in the marked diversity of these cellular forms emerging from a single replicative unit. It was unknown how JCVI-syn3.0 propagated in the absence of controlled FtsZ-based cell division, as in JCVI-syn1.0 and JCVI-syn3A. We observed that genomically minimized strains displayed a wide variety of dynamics (Figs 1F, 3, 4E). Filamentous growth was notable, in particular, given the lack of a cell wall or bacterial cytoskeletal elements to generate and stabilize these cellular forms. While some filaments exhibited branching or pearling, we did not observe scission, consistent with the presence of an energy barrier (Beltrán-Heredia et al., 2017; Caspi and Dekker, 2014; Ruiz-Herrero et al., 2019).

These growth characteristics appeared similar to L-form bacteria, which are variants that lack their typical cell wall (Errington, 2017). Mycoplasmas and L-forms have long been compared to one another, as both lack a cell wall and appear similar in optical (Kang and Casida, 1967) and electron micrographs (Dienes and Bullivant, 1968). Given these morphological similarities, genomically minimized mycoplasmas and L-forms may share common mechanisms of propagation. In particular, some L-forms undergo an irregular cell division mechanism, termed “extrusion-resolution,” involving extrusion of excess membrane, followed by resolution of the extrusion into connected units (Errington, 2017). This mechanism requires excess membrane synthesis (Mercier et al., 2013), does not require FtsZ or FtsA (Leaver et al., 2009), and is suppressed by confinement in submicron microfluidic channels (Wu et al., 2020). Our observations suggest the extrusion-resolution model is relevant to propagation of JCVI-syn3.0 and related genomically minimized strains, and our study highlights a wide range of morphological dynamics that can occur without shear flow.

### Systematic approach attributes morphology to genes of unknown function

We leveraged our modular genome design to determine a set of seven genes, all of which were required together to recover a more normal morphology in a genomically minimal context. In L-forms, restoration of cell wall synthesis can recover normal morphology (Kawai et al., 2014), but normal morphology has not to our knowledge been recovered in L-forms still lacking a cell wall, unlike in the mycoplasmas here. The viability of L-forms depends on multiple classes of mutations affecting peptidoglycan precursor synthesis, fatty acid synthesis, glycolysis, or oxidative stress response (Errington, 2017). Due to the complexity of such a phenotype, we therefore took an approach agnostic to gene function and first screened multiple genome segment variants, followed by gene clusters, and finally individual genes.

This systematic approach enabled the identification of proteins with previously unknown function as necessary for normal morphology in the minimal cell. Although their functions are unclear, genes *527* and *604* are expressed abundantly in JCVI-syn3A, with several hundred copies per cell, whereas others (genes *520, 521, 605*) were detected in lower quantities (Breuer et al., 2019). Furthermore, some protein sequence motifs are highly conserved across bacterial species. Gene *0527* is homologous to the DUF177 family, which may participate in biosynthesis of membrane proteins (Yang et al., 2016), while gene *604* is homologous to LemA/GacS, a family of two-component regulatory proteins (Hrabak and Willis, 1992). Two-component regulatory systems have not been reported in mycoplasmas (Capra and Laub, 2012), nor are any two-component genes other than the *lemA*/*gacS* gene observed here, leaving the function of gene *604* unclear in JCVI-syn3A. Secondary structural analyses further revealed that some of these unknown proteins are associated with the membrane, including genes *604* and *605* that encode, respectively, one transmembrane helix and a bacterial lipoprotein anchoring motif. Gene *602* is annotated as a pseudogene, due to a confirmed frameshift mutation causing truncation relative to full-length orthologs in the *Mycoplasma mycoides* group of organisms. The N-terminal portion, predicted to be expressed, contains two transmembrane helices. Therefore, while these four genes outside the *dcw* cluster lack known function, bioinformatic analyses indicate they associate with the membrane, suggesting unknown roles for membrane properties in cell morphology and division.

### JCVI-syn3A highlights aspects of Z-ring function

Despite extensive characterization of the multicomponent systems underlying cell division in walled bacteria, a full understanding of cell propagation in those organisms is far from complete (Margolin, 2020; Osawa and Erickson, 2018). Gaps in understanding include spatiotemporal control and interactions among the myriad proteins involved, as well as mechanisms of force transduction. In phylogenetically related mycoplasmas in the *M. mycoides* group, only incomplete models of cell division have been reported (Seto and Miyata, 1998). As a framework to simplify and study the genetic basis of cell division, JCVI-syn1.0 already lacks most cell division components of more complex bacteria.

JCVI-syn3.0 and JCVI-syn3A present compelling platforms to study genetic requirements for cell division and morphological control, as many alterations to the genotype resulted in readily observable phenotypic variation. In particular, FtsZ was required but alone insufficient for cell division in JCVI-syn3.0+126, evidencing contributions of other proteins to cell division. JCVI-syn3.0+126 retains FtsA and SepF, which can anchor FtsZ to the membrane (Duman et al., 2013; Hamoen et al., 2006; Loose and Mitchison, 2014), as well as modulate the assembly and bundling of FtsZ (Krupka and Margolin, 2018; Singh et al., 2008). FtsA can polymerize *in vitro* (Szwedziak et al., 2012), but the proteome of JCVI-syn3A includes only ≈40 copies per cell, which is significantly less abundant than FtsZ at ≈640 copies per cell (Breuer et al., 2019). In both *Escherichia coli* and *Bacillus subtilis*, cell division occurs when FtsZ accumulates to a threshold number per cell, although its molecular mechanism and range of interactions with other proteins remain unclear (Sekar et al., 2018; Si et al., 2019). Other *dcw* cluster genes, *mraZ* and *mraW*, are retained in JCVI-syn3.0. However, recent transposon mutagenesis suggests these are not essential for growth, and MraZ was not detected in the JCVI-syn3A proteome (Breuer et al., 2019). JCVI-syn3A lacks genes known to facilitate other aspects of Z-ring function, such as FtsZ degradation and positioning. Because purified FtsZ protofilaments are intrinsically curved (Housman et al., 2016), we speculate that alignment of FtsZ along maximal membrane curvature may help localize FtsZ to the division furrow, similar to the orientation of another intrinsically curved filament, MreB, along the maximal membrane curvature in *B. subtilis* (Hussain et al., 2018).

### Minimal cells highlight membrane biophysics in cell division

The complex role of FtsZ, as well as the requirement for genes of unknown function, indicate the contribution of other processes to controlled cell division and maintenance of normal morphology in JCVI-syn3A and JCVI-syn3.0+126. Both our imaging and genetics results indicate the influence of membrane properties on cell morphology. In microfluidic chemostats, JCVI-syn3.0 and RGD6 displayed aberrant membrane-bound vesicles, consistent with dysregulation of membrane accumulation linked with pleomorphism. Bioinformatic analyses suggest membrane association of the products of the four required genes of unknown function outside the *dcw* cluster. Maintenance of membrane homeostasis is critical in cellular organisms, including bacteria (Ernst et al., 2016; Zhang and Rock, 2008), but is not well understood in mycoplasmas, which have adapted through evolution to acquire membrane components directly from the environment (Breuer et al., 2019). Membrane components from the environment can affect mycoplasmal morphology. For example, the lipid composition of the growth medium affects whether mycoplasmas phylogenetically similar to JCVI-syn1.0 form spherical cells or filaments (Razin et al., 1967). Furthermore, cell division requires membrane synthesis in mycoplasmas phylogenetically similar to JCVI-syn1.0 (Seto and Miyata, 1999). Theories and observations of abiotic vesicles predict a wide range of morphological dynamics similar to those observed here, including spontaneous filamentation and pearling (Caspi and Dekker, 2014; Lipowsky, 2013), which depend on membrane properties, such as fluidity, elastic modulus, and spontaneous curvature. With a defined, nearly minimal genome and an experimental niche that can be manipulated to deliver lipids, JCVI-syn3A presents an ideal model system for examining the role of membrane dynamics in cellular propagation.

### Conclusions

We present here the first use of genomically minimized cells to determine the genetic basis for the core physiological processes of cell division and the maintenance of cell morphology. Starting with the minimal cell JCVI-syn3.0, which shows uncontrolled cell division and pleomorphism, we reconstituted a set of genes that conferred nearly normal division and morphology. The resulting strain, JCVI-syn3A, has a nearly minimal genome and metabolism, as well as cell morphology typical of most spherical bacteria. JCVI-syn3A thus offers a compelling minimal model for bacterial physiology and platform for engineering biology broadly. Of the 19 additional genes present in JCVI-syn3A, seven were required and together sufficient to restore the normal phenotype. Five of these genes have no known function. Our systematic approach, agnostic to gene function, discovered their requirement for normal cell division and morphology and may find application to organisms beyond mycoplasmas. The role of these previously uncharacterized genes, and the polygenic basis of the phenotype, will inform bottom-up approaches to reconstitute cell division in synthetic cells.

## Author contributions

JFP, LS, KSW, JIG, and EAS authored the manuscript. JFP and EAS fabricated the microfluidic chemostats and performed microfluidic measurements. LS performed experiments to reintroduce genes and identify those resulting in improved morphological uniformity of JCVI-syn3.0. LS and KSW provided optical microscopy of wet mounts. LS, NAG, BJK, and RYC generated the bacterial strains. TJD and MHE performed electron microscopy. AM and NG assisted in developing the microfluidic device and its operation.

## Acknowledgements

JFP was supported by a Fannie and John Hertz Foundation Fellowship, the Mitchison lab at Harvard Medical School, and the Fakhri lab at MIT. EAS was supported by the National Institute of Standards and Technology: certain commercial equipment, instruments, or materials are identified in this paper to foster understanding; such identification does not imply recommendation or endorsement by the National Institute of Standards and Technology, nor does it imply that the materials or equipment identified are necessarily the best available for the stated purpose. LS, KSW, and JIG were supported by National Science Foundation grants MCB 1840320 and MCB 1818344 to JIG. LS, KSW, NAG, BJK, RYC, and JIG were supported by Synthetic Genomics, Inc., the Defense Advanced Research Projects Agency’s Living Foundries program (Contract HR0011-12-C-0063), and the J. Craig Venter Institute. Microscopy work at the University of California**-**San Diego was supported by NIH grant P41GM103412 from the National Institute of General Medical Sciences to MHE.

## Methods

- **Fig S1**. Explanation of microfluidic chemostats
- **Fig S2**. Location of cytoplasmic protein, nucleoids, and membrane in pleomorphic cells
- **Fig S3**. Overview of genomic manipulations to generate strains reported in this study
- **Fig S4**. Images of strains to determine genes required for normal morphology
- **Table S1**. Overview of genomically minimized strains and derivatives
- **Table S2**. Primers used in this study
- **Table S3**. Expected sizes of amplicons using junction primers for strains
- **Table S4**. Strains constructed to determine genes required for normal morphology
- **SI Movie 1. Growth of JCVI-syn1.0 in microfluidic chemostats**. Related to Fig 1E. Cells were submicron and largely disconnected from one another. Top panel: Phase contrast. Bottom panel: Constitutively expressed mCherry, as a marker for cytoplasmic protein. Scale bar: 5 µm.
- **SI Movie 2. Growth of JCVI-syn3.0**. Related to Fig 1F. Some cells were several microns in diameter or had irregular shapes, including filamentous cellular forms. Individual cells exhibited dynamic morphological transitions resulting from processes intrinsic to the cells. Top panel: Phase contrast. Bottom panel: Constitutively expressed mCherry, as a marker for cytoplasmic protein. Scale bar: 5 µm.
- **SI Movie 3. Fluctuations of cells in a microfluidic chemostat**. Related to Fig 1. Correlated fluctuations of adjacent substructures suggest physical connections between them. Fluctuations of cells in the microfluidic chemostat also indicate sufficient passivation of the chemostat surfaces.
- **SI Movie 4. Growth of RGD6, including a long filamentous cell**. Related to Fig 3A. A single RGD6 grew into a filament in the absence of shear flow, due to morphological dynamics intrinsic to the cell. Vesicles lacking constitutively expressed mCherry initiated at the ends of the filament and increased in size. Top panel: Phase contrast. Bottom panel: Constitutively expressed mCherry, as a marker for cytoplasmic protein. Scale bar: 5 µm.
- **SI Movie 5. Growth of RGD6, including a large vesicle**. Related to Fig 4E. A large vesicle lacking constitutively expressed mCherry increased in size. Cells containing mCherry were contiguous with the surface of the vesicle. Top panel: Phase contrast. Bottom panel: Constitutively expressed mCherry, as a marker for cytoplasmic protein. Scale bar: 5 µm.
- **SI Movie 6. Growth of JCVI-syn3A**. Related to Fig 4F. JCVI-syn3A is a nearly minimal cell with 19 more genes than JCVI-syn3.0 and exhibits a nearly normal morphology similar to JCVI-syn1.0. Scale bar: 5 µm.

### Synthetic M. mycoides strains, growth media

The principal mycoplasmal strains used in this study are JCVI-syn1.0 (genome sequence CP002027), JCVI-syn3.0 (genome sequence CP014940.1), and JCVI-syn3A (genome sequence CP016816.2). Chromosomes of all strains and their derivatives carry a yeast *CEN*/*ARS* and a *his3* marker for centromeric propagation of the genome in yeast, as well as a *tetM* resistance marker for selection after genome transplantation into a mycoplasma recipient. Mycoplasmal strains were propagated in SP4 medium containing fetal bovine serum (hereafter SP4), as previously described (Hutchison et al., 2016).

### Microscopy of bulk cultures

Mycoplasma transplants were grown in static, planktonic culture at 37 °C in SP4 liquid medium without tetracycline. To observe cell morphologies in culture with minimal manipulation, wet mounts were prepared from logarithmic phase cultures after 3 days of growth, by depositing 3 µL of settled cells, which had been carefully removed by micro pipette tip from round-bottom culture tubes, onto an untreated glass slide and applying an 18 × 18 mm coverslip. Optical microscopy was performed using a Zeiss Axio Imager 1 microscope with a Zeiss plan/apochromatic 63x oil 1.4 objective, differential interference contrast (DIC) optics and X-Cite 120PC Q mercury arc lamp. A 43HE filter with excitation BP 550/25 (HE) and emission BP 605/70 (HE) was used to detect fluorescence in constructs containing mCherry.

### Empirical gradient thresholding to estimate cell size distributions

Size distributions were quantified using the empirical gradient threshold (EGT) algorithm to binarize grayscale images (Chalfoun et al., 2015).

### Scanning electron imaging

Cells grown in SP4 medium were centrifuged at room temperature for 4 min at 2000 g to produce a loose pellet. 1 mL of fixative solution consisting of 2.5 % (w/v) glutaraldehyde, 100 mmol/L sodium cacodylate, 2 mmol/L calcium chloride, and 2 % (w/v) sucrose was added cold and replaced 950 µL of growth medium. Samples were subsequently stored at 4 °C. Imaging substrates were glass coverslips coated with indium tin oxide and treated with polyethylenimine or poly-D-lysine. (10 to 20) µL of solution containing cells were placed on top of the coverslips for 2 min and subsequently washed five times for 2 min each on ice using 0.1 mol/L cacodylate buffer with 2 mmol/L calcium chloride and 2 % (w/v) sucrose. Cells were fixed further on ice in 2 % (w/v) osmium tetroxide with 2 % (w/v) sucrose in 0.1 mol/L cacodylate for 30 min, followed by rinsing with double distilled water and dehydration in an ethanol series of (20, 50, 70, and 100) % (v/v) for 2 min each on ice. Substrates with immobilized cells were then dried through the critical point with carbon dioxide and sputter-coated with a thin layer of Au/Pd. Images were collected with a Zeiss Merlin Fe-SEM at 2.5 keV, 83 pA probe current, and 2.9 mm working distance using zero tilt and the secondary electron detector.

### Microfluidic platform fabrication

We fabricated microfluidic chemostats to culture and image the growth of genomically minimized mycoplasmas in a shear-free, biochemically controlled environment. Shallow growth chambers confined cells to facilitate imaging, while deeper microfluidic channels loaded cells, perfused fresh growth media, and introduced fluorescent labels to determine the spatial distribution of cytoplasmic protein, nucleoids, membrane, and extracellular growth medium. The concept of the device is similar to previous microfluidic chemostats (Wang et al., 2010), with the exception cells in our chambers were free to diffuse.

Soft lithographic processing allowed facile fabrication and characterization of devices compatible with cell growth and optical imaging. A negative master was fabricated on a silicon wafer, using reactive ion etching to define 3.1 µm deep chambers arrayed along a 100 µm wide by 22 µm deep channel defined by SU-8 photoresist (MicroChem). After treating with a silane release layer, a positive mold of the device was created in polydimethylsiloxane (PDMS, Sylgard 184) and bonded irreversibly to a #1.5 borosilicate coverslip after treatment with an oxygen plasma. Correcting for known 1.4 % shrinkage after curing at 80 °C, chamber and channel depths decrease to 3.0 µm and 21 µm, respectively. The lateral dimensions of the chambers ensured rapid diffusive exchange with the deeper channel and are visible in Figs 1, 3, 4, S1, and S2, and SI Movies. Devices were filled with fluid immediately, to take advantage of the small liquid contact angle after plasma treatment.

**Figure S1.**
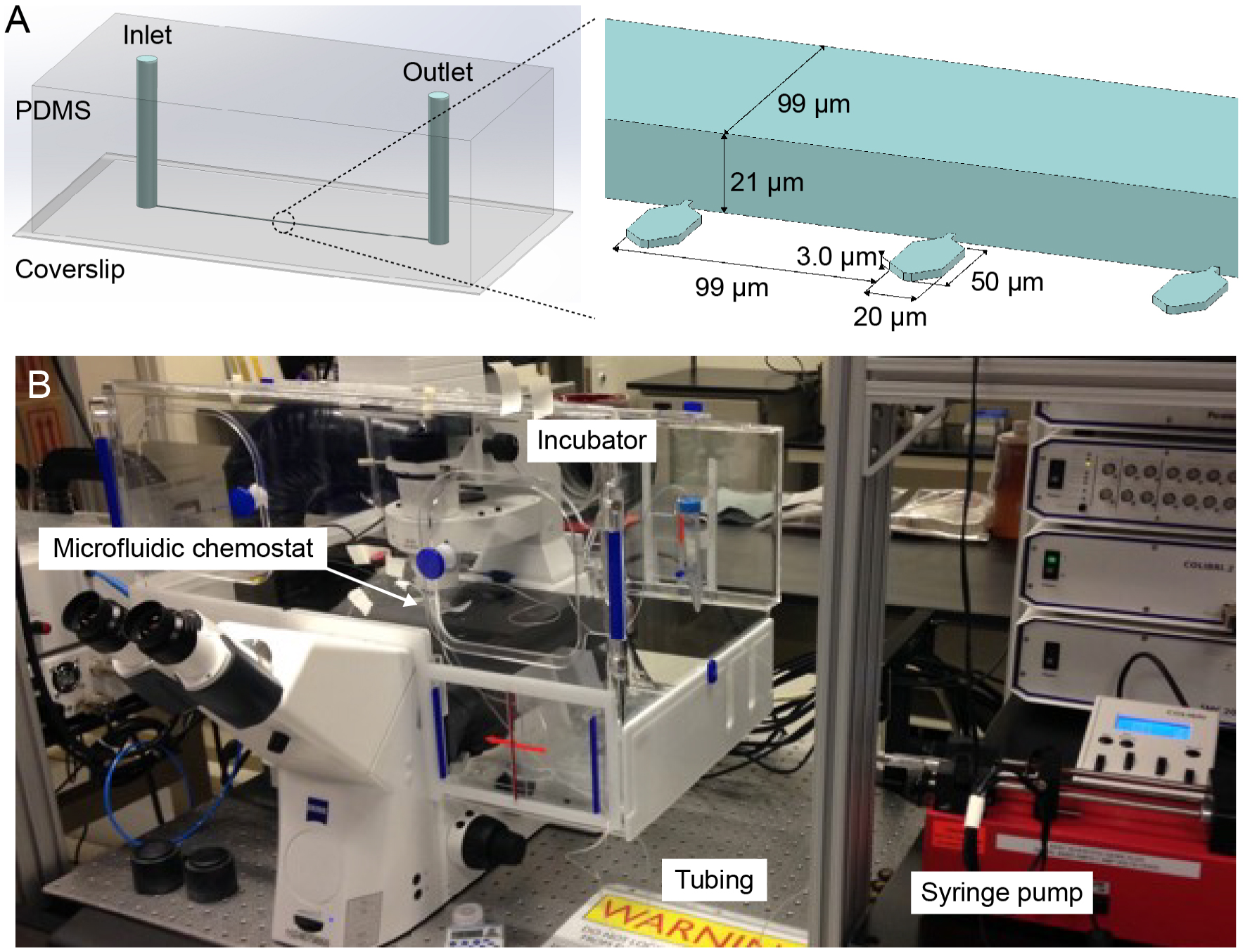
Microfluidic chemostat isolates cells from shear flow to image their intrinsic morphological dynamics during growth and propagation. Related to Figures 1, 3, and 4. (A) The microfluidic chemostat is shown schematically with internal dimensions. Cells grow in chambers, in diffusive contact with a flow channel that provides a continuous supply of fresh growth medium. (B) Microfluidic chemostat shown within the complete laboratory setup that includes the syringe pump and incubator.

### Microfluidic cell culture and microfluidic optical imaging

Cells were imaged using an inverted microscope outfitted with an incubator and heated stage, set to 37.0 °C, vibration isolation workstation, 100x magnification objective (Zeiss, alpha Plan-Apochromat, 1.46 numerical aperture, oil immersion), suitable LED excitation sources and emission filter cubes, and a water-cooled sCMOS camera (Hamamatsu Orca Flash 4.0 v2). Cells were grown in SP4 medium, but with the pH indicator dye phenol red omitted to reduce background fluorescence. Image acquisition was automated using Zeiss Zen software and automated focus correction, for multichannel, timelapse imaging of overnight growth experiments over ≈(10 to 16) h, with images taken every ≈(10 to 30) min, as well as endpoint staining with fluorescent dyes. As appropriate to each strain, cells were imaged overnight in transilluminated brightfield and fluorescence, to image the cytoplasm of cells expressing mCherry. Endpoint fluorescence labels included 10 µg/mL Hoechst 33258 (Life Technologies) to visualize DNA, 84 µg/mL SP-DiOC18(3) (Life Technologies) to visualize membranes, and FITC-Dextran (10 kDa molecular weight) to image the negative space around cells. Cells were incubated with the fluorescent dyes for ≈(5 to 100) min in SP4 medium or in PBS or Tris sucrose buffer (10 mmol/L Tris pH 6.5 and 0.5 mol/L sucrose) to reduce background fluorescence. Strains expressing mCherry exhibited qualitatively similar morphological features as strains without mCherry, so mCherry strains were useful to investigate the composition of different morphotypes.

Typically, device surfaces were functionalized with 0.1 mg/mL poly-L-lysine-g-polyethylene glycol (SuSoS) in 10 mmol/L HEPES pH 7.4 soon after bonding, to prevent cell adhesion. After incubating for 40 min with PLL-g-PEG, devices were flushed with sterile water, followed by SP4 growth medium and cells in the same growth medium. For microfluidic cell culture, SP4 medium was prepared without phenol red, to reduce background signal during fluorescence imaging. Cells entered the chambers diffusively at room temperature for ≈(5 to 60) min, depending on the typical cell size, cell concentration in the loading channel, and the desired number of cells per chamber. Devices mounted on a microscope for cell growth and imaging were infused with fresh growth medium at 200 µL/hr using a syringe pump.

The small submicron cells diffused freely in the chambers but tended to congregate opposite the chamber entrance, likely due to a small net fluid flow into the bulk device material (Kolnik et al., 2012; Randall and Doyle, 2005). Thus, in contrast to related mother machine devices (Wang et al., 2010), the number of cells per chamber increased over time.

**Figure S2.**
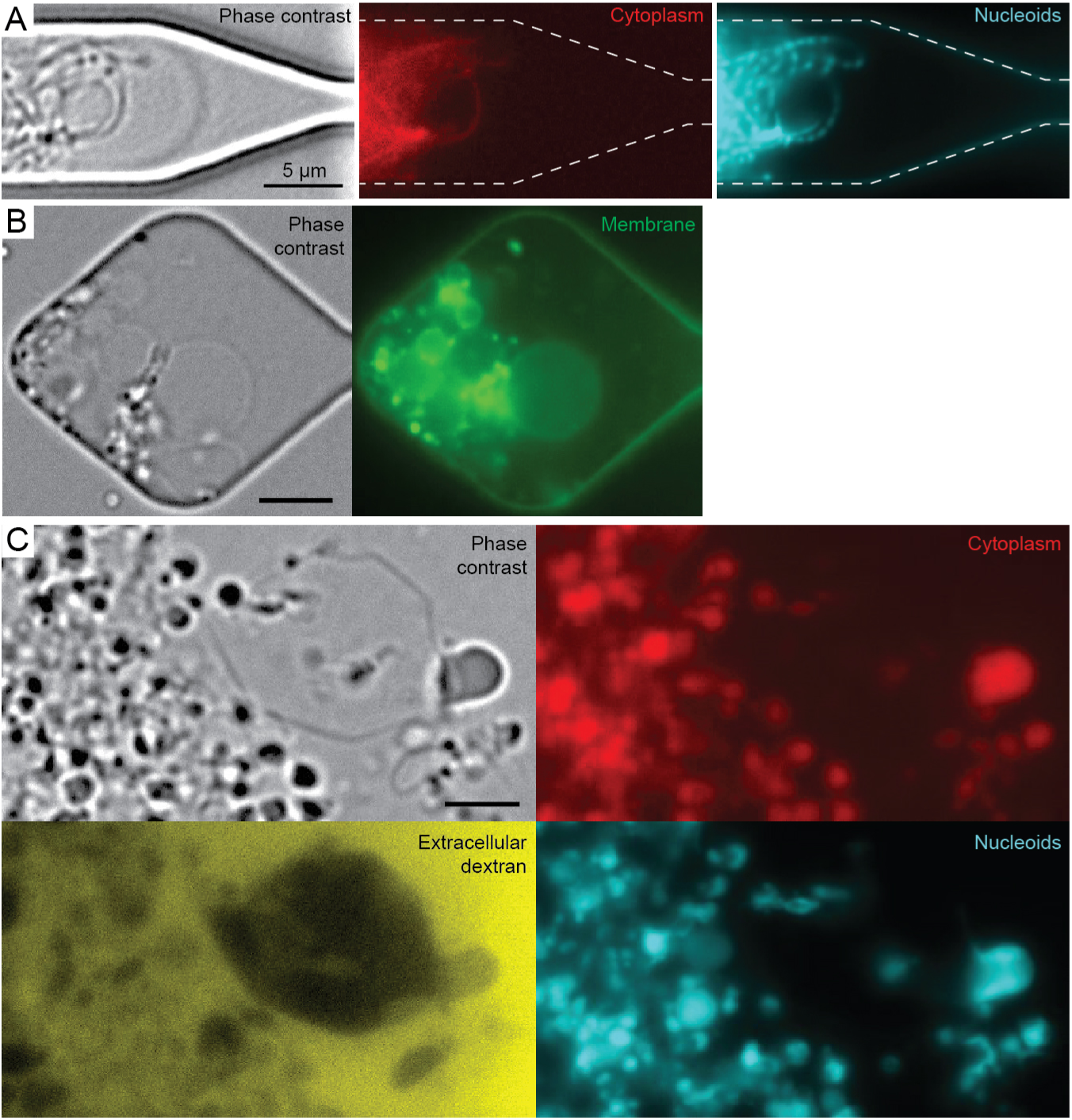
Location of cytoplasmic protein, nucleoids, and membrane in pleomorphic cellular forms. Brightfield and fluorescence optical micrographs of RGD6+mCherry grown in microfluidic chambers show filamentous cells (A), and large cells and vesicles (B,C). (A) Nucleoids appeared separated along the length of filamentous cells, suggesting genome segregation may continue in the absence of complete cell scission, as recently observed in *B. subtilis* L-forms (Wu et al., 2020). (B) The surface of vesicles, which appeared as lower contrast in phase contrast images, were stained with the membrane dye SP-DiOC18(3). (C) A large vesicle lacked constitutively expressed mCherry but excluded FITC-conjugated dextran in the growth medium, suggesting the vesicle membrane was not permeable to macromolecules. Scale bars: 5 µm (A,B) and 2 µm (C).

**Table S1.**
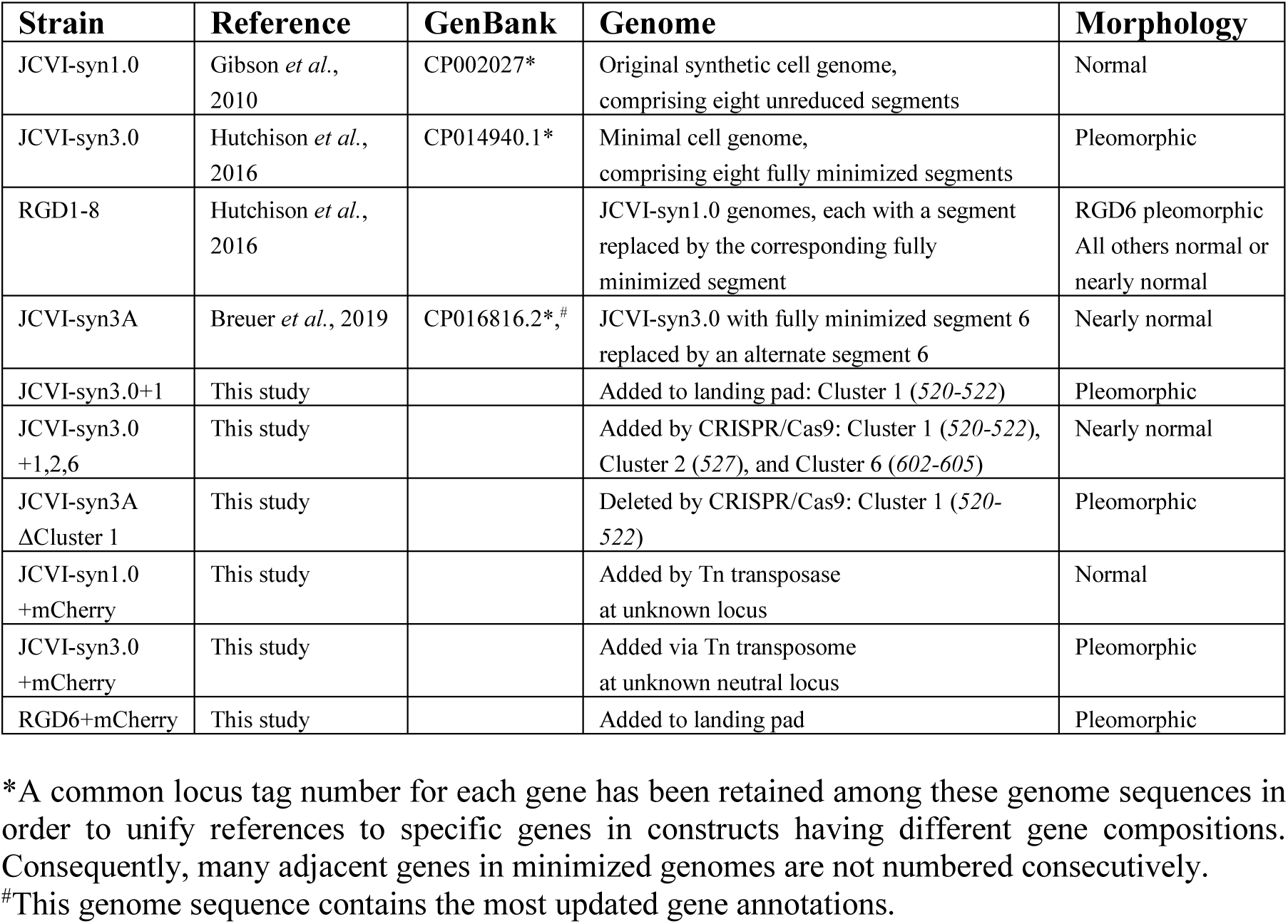
Overview of genomically minimized strains and derivatives.

| Strain | Reference | GenBank | Genome | Morphology |
| --- | --- | --- | --- | --- |
| JCVI-syn1.0 | Gibson *et al.*, 2010 | CP002027* | Original synthetic cell genome, comprising eight unreduced segments | Normal |
| JCVI-syn3.0 | Hutchison *et al.*, 2016 | CP014940.1* | Minimal cell genome, comprising eight fully minimized segments | Pleomorphic |
| RGD1-8 | Hutchison *et al.*, 2016 |  | JCVI-syn1.0 genomes, each with a segment replaced by the corresponding fully minimized segment | RGD6 pleomorphic<br>All others normal or nearly normal |
| JCVI-syn3A | Breuer *et al.*, 2019 | CP016816.2*,# | JCVI-syn3.0 with fully minimized segment 6 replaced by an alternate segment 6 | Nearly normal |
| JCVI-syn3.0+1 | This study |  | Added to landing pad: Cluster 1 (520-522) | Pleomorphic |
| JCVI-syn3.0+1,2,6 | This study |  | Added by CRISPR/Cas9: Cluster 1 (520-522), Cluster 2 (527), and Cluster 6 (602-605) | Nearly normal |
| JCVI-syn3A ΔCluster 1 | This study |  | Deleted by CRISPR/Cas9: Cluster 1 (520-522) | Pleomorphic |
| JCVI-syn1.0 +mCherry | This study |  | Added by Tn transposase at unknown locus | Normal |
| JCVI-syn3.0 +mCherry | This study |  | Added via Tn transposome at unknown neutral locus | Pleomorphic |
| RGD6+mCherry | This study |  | Added to landing pad | Pleomorphic |

\*A common locus tag number for each gene has been retained among these genome sequences in order to unify references to specific genes in constructs having different gene compositions. Consequently, many adjacent genes in minimized genomes are not numbered consecutively.
#This genome sequence contains the most updated gene annotations.

**Figure S3.**
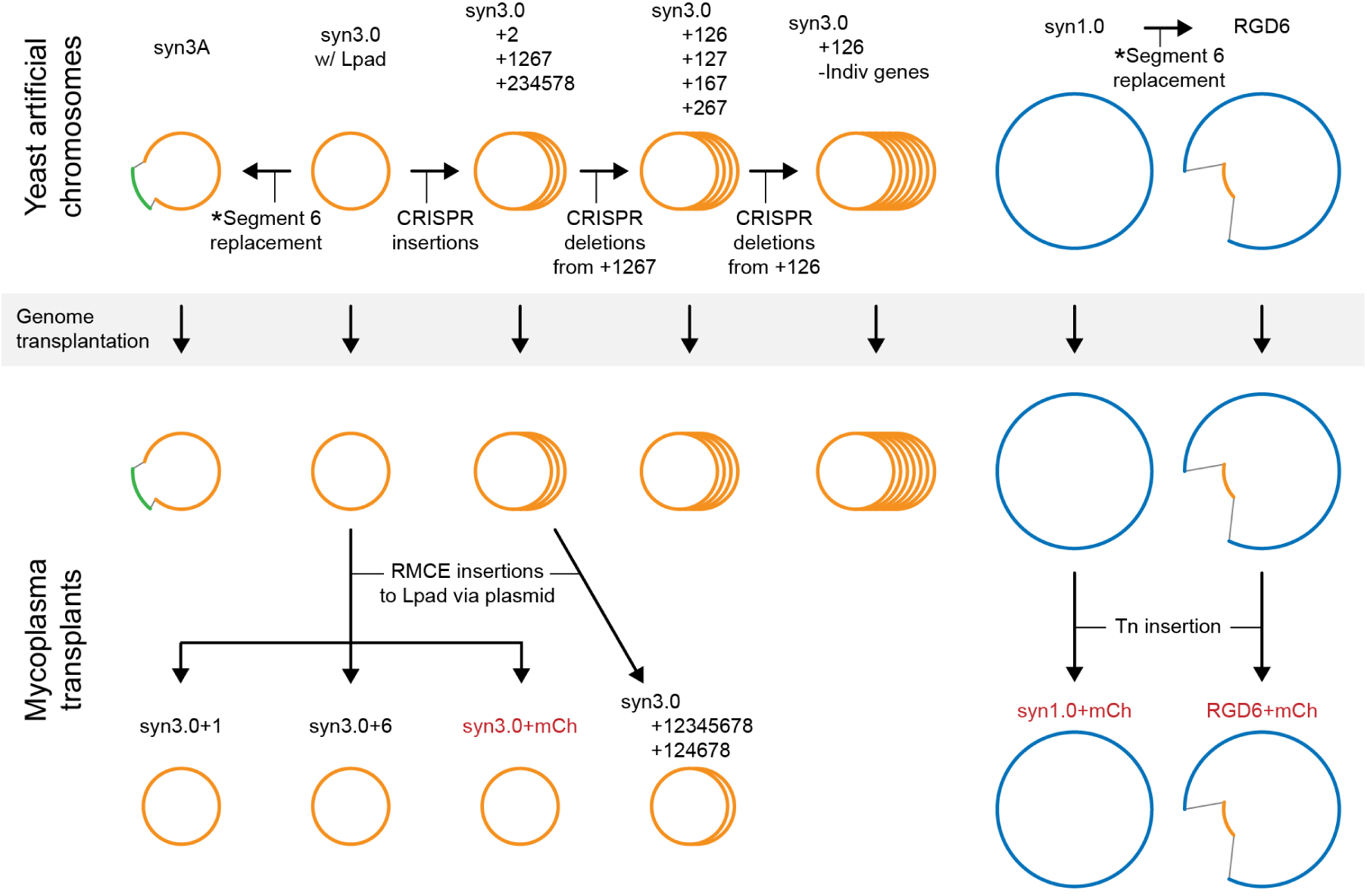
Overview of genomic manipulations to generate strains reported in this study. Genomes were manipulated using two strategies: (1) For artificial chromosomes in yeast (upper constructs) CRISPR/Cas9 (CRISPR) methodology was used to generate insertions/deletions, followed by genome transplantation into a mycoplasma recipient to render mycoplasmal organisms programmed with the new genome; or, (2) for chromosomes in the mycoplasmal transplants (lower constructs), recombination-mediated-cassette-exchange (RMCE) was used to introduce genes transferred from plasmids to a chromosomal landing pad (Lpad) comprising heterospecific *loxP* sites. In cases where the landing pad was absent (JCVI-syn1.0 and RGD6), a gene encoding the fluorescent protein mCherry was introduced by transposome mediated insertion. The key gene clusters transferred, numbered 1 through 8, correspond to those indicated in Figure 5 of the article. The content of gene clusters within each strain or group of strains is indicated according to the designations in Figure 5. A group of strains (tiled circles) with different combinations is indicated by listing the combinations represented. Arrows represent a manipulation from one strain to another. Arrows in the shaded area indicate genome transplantations. Details of these processes are described in the following supplementary information. *Methods for replacement of genomic segments, as well as the construct RGD6, are detailed elsewhere (Hutchison et al., 2016). The full sequence and annotation of segment 6 in JCVI-syn3A is provided in the whole genome sequence (GenBank accession CP016816.2).

### Genome synthesis and assembly

The synthesis and assembly of minimized genomes has been described previously (Hutchison et al., 2016). Briefly, overlapping oligonucleotides of < 80 bases were designed using WordPerfect macros (Hutchison et al., 2016). Streamlined oligonucleotide design software was also embedded in the Archetype® software available commercially through Codex DNA, Inc. Then, oligonucleotides were purchased (Integrated DNA Technologies), pooled, assembled into 1.4 kilobase fragments, error corrected, and amplified. These fragments were subsequently assembled into 7 kilobase cassettes using Gibson Assembly and cloned in *E. coli*. These cassettes were further assembled into genome segments and whole genomes in *S. cerevisiae*.

### Construction of genomes by combining genome segments

The final assembly of whole genomes by combining chromosome segments has been described previously (Hutchison et al., 2016). The complete genome sequences of JCVI-syn1.0 and the minimal JCVI-syn3.0, comprised of a combination of each minimal segment 1 through 8, are available through GenBank (Table S1). Additional constructs were made (Hutchison et al., 2016) by placing each of the eight fully-minimized segments into a genome containing the seven other unminimized segments of the JCVI-syn1.0 genome. One resulting strain contains a minimal segment 6 in an otherwise unminimized genome (RGD6) and is further characterized here. Another construct, JCVI-syn3A, comprises the minimal JCVI-syn3.0 genome whose segment 6 is replaced with an alternate version that includes 19 additional genes. The complete genome sequence of this construct is available through GenBank (Table S1) and represents the most recent and accurate annotation of genes.

### Gene additions to JCVI-syn3.0 via Cre-Lox recombination and plasmid-mediated cassette exchange (RMCE)

JCVI-syn3.0 subclone 13-2, referred to simply as JCVI-syn3.0 in this study, contains two hetero-specific *loxP* sites that comprise a landing pad between genes *601* and *606*, which are adjacent in the fully-minimized JCVI-syn3.0 genome (Hutchison et al., 2016; Noskov et al., 2015). Plasmid Pmod2-loxpurolox-sp-cre (annotated sequence in GenBank: MN982903.1) includes the puromycin resistance marker flanked by two hetero-specific *loxP* sites and serves as a vector to transfer gene cassettes into JCVI-syn3.0 using Cre recombinase/*loxP*-mediated recombination. After transformation into recipient JCVI-syn3.0 cells, the *loxP*-flanked cassette in the plasmid inserts into the genome, replacing the *loxP*-flanked region in the genome.

### Protocol for transformation of JCVI-syn3.0 using plasmids

Cells were transformed as described previously (Hutchison et al., 2016). Briefly, JCVI-syn3.0 cells were grown in 4 mL of SP4 growth medium to reach pH 6.5 to pH 7.0. The culture was centrifuged for 15 min at 4700 rpm and 10 °C in a 50 mL centrifuge tube. The pellet was resuspended in 3 mL of Sucrose/Tris (S/T) Buffer, composed of 0.5 mol/L sucrose and 10 mmol/L Tris at pH 6.5. The resuspended cells were centrifuged for 15 min at 5369 g and 10 °C. The supernatant was discarded, and the pellet resuspended in 250 µL of 0.1 mol/L CaCl2 and incubated for 30 min on ice. Then, 200 ng of plasmid was added to the cells, and the centrifuge tube was mixed gently. 2 mL of 70 % (w/v) polyethylene glycol (PEG) 6000 (Sigma), dissolved in S/T Buffer, was added to the centrifuge tube, and mixed well using a serological pipette. After a 2 min incubation at room temperature, 20 mL S/T Buffer without PEG was added immediately and mixed well. The tube was centrifuged for 15 min at 10,000 g and 8 °C. The supernatant was discarded, and the tube inverted with the cap removed on tissue paper to drain residual PEG. The cells were subsequently resuspended in 1 mL of SP4 growth medium prewarmed to 37 °C. These cells were incubated for 2 h at 37 °C, followed by plating on SP4 agar containing 3 µg/mL of puromycin (Sigma). Colonies appeared after 3 to 4 days at 37 °C.

### Construction of plasmid Pmod2-loxpurolox-sp-cre_520-522 (cluster 1)

The plasmid Pmod2-loxpurolox-sp-cre_*520-522* was constructed by inserting the genes *520-522* adjacent to the puromycin resistance marker in plasmid Pmod2-loxpurolox-sp-cre via assembly as previously described (Kostylev et al., 2015). The genome cassette spanning genes *520-522* was amplified with primers M_520_F and M_520_R, PrimeSTAR Max Premix (Takara), and using JCVI-syn1.0 genomic DNA as the template. PCR conditions were: 98 °C for 2 min; 30 cycles of 98 °C for 10 s, 55 °C for 10 s, and 72 °C for 1 min; and, 72 °C for 2 min. The linear vector Pmod2-loxpurolox-sp-cre was amplified using primers M_v_F and M_v_R, PrimeSTAR Max Premix, and using the plasmid Pmod2-loxpurolox-sp-cre as the template. PCR conditions were: 98 °C for 2 min; 30 cycles of 98 °C for 10 s, 58 °C for 10 s, and 72 °C for 1 min; and, 72 °C for 2 min. The PCR product for the linear vector was mixed with 1 µL of DpnI and incubated at 37 °C for 1 h to digest the plasmid template. The PCR amplicon was purified using a DNA Clean & Concentrator Kit (Zymo Research). Purified DNA fragment *520-522* and linear vector were introduced by transformation into competent cell DH5alpha C2987H (New England Biolabs). 1 ml of SOC (Thermo Fisher Scientific) was added to the transformation mix, and, after incubation at 37 °C for 1 h, cells were plated on AmpR LB agar plates and incubated at 37 °C overnight. Colony PCR was used to screen surviving transformants for the presence of the *520-522* insert using primers spanning the insert junctions and amplification with Taq 2X Master Mix PCR Kit (New England Biolabs), as described by the supplier. The PCR conditions were: 94 °C for 3 min; 30 cycles of 94 °C for 15 s, 55 °C for 20 s, and 68 °C for 30 s; and, followed by 68 °C for 1 min. The expected DNA sizes are listed in Table S3. Positive clones with the predicted 330 bp DNA band indicating the insertion of genes *520-522* were identified. Plasmids from positive clones were extracted using a Qiagen Mini-Prep Kit by the manual. Sanger sequencing of plasmids confirmed the absence of mutations. Primers for PCR amplification, junction colony PCR, and sequencing are listed in Table S2.

### Construction of plasmid Pmod2-loxpurolox-sp-cre_602-605 (cluster 6)

The plasmid Pmod2-loxpurolox-sp-cre_*602-605* was constructed by inserting the genes *602-605* adjacent to the puromycin resistance marker in Pmod2-loxpurolox-sp-cre via Gibson Assembly. The genes *602-605* were amplified using primers 0602_fwd and 0605_rev, PrimeSTAR Max Premix, and JCVI-syn1.0 genome as the template. The linear vector Pmod2-loxpurolox-sp-cre was PCR amplified using primers vec_fwd and vec_rev, PrimeSTAR Max Premix, and the plasmid Pmod2-loxpurolox-sp-cre as the template. The PCR conditions were: 98 °C for 2 min; 30 cycles of 98 °C for 10 s, 55 °C for 20 s, and 72 °C for 40 s; and, followed by 72 °C for 5 min. Zymo DNA Clean & Concentrator Kit purified DNA fragment *602-605* and gel purified linear vector were assembled via Gibson Assembly. The assembly mix was transformed using competent cell DH5alpha C2987H, grown at 37 °C for 1 h and plated on an AmpR LB plate. The plate was incubated at 37 °C overnight. Colony PCR was used to screen positive colonies with genes *602-605* insertion in vector using junction primers and Qiagen Multiplex PCR Kit by the manual. The expected DNA sizes are listed in Table S3. Positive clones showing the 335 bp and 507 bp PCR products that confirmed genes *602-605* insertion were obtained. Plasmids from positive clones were extracted using a Qiagen Mini-Prep Kit. Sanger sequencing of the plasmids confirmed the absence of mutations. Primers for PCR amplification, colony PCR, and sequencing are listed in Table S2.

### Confirmation of sequences in transformants

To confirm that the genes *520-522* or *602-605* and the puromycin resistance marker were inserted into the JCVI-syn3.0 genome after transformation, a diagnostic PCR with PrimeSTAR Max Premix was performed to screen positive colonies with the complete JCVI-syn3.0 genome, the genes *520-522* or *602-605*, and the puromycin resistance marker using transformant junction primers. For the cluster *520-522*, primers 0520_L(486) and pgk_R(486) were used. The PCR conditions were: 98 °C for 2 min; 30 cycles of 98 °C for 10 s, 55 °C for 20 s, and 72 °C for 30 s; and, followed by 72 °C for 2 min. For cluster *602-605*, primers seq_F1 and puro_R were used. The PCR conditions were: 98 °C for 2 min; 30 cycles of 98 °C for 5 s, 55 °C for 10 s, and 72 °C for 40 s; and, followed by 72 °C for 5 min.

The genomic DNA of positive colonies was extracted, and regions *520-522* or *602-605* were amplified via PCR with primers seq_F1 and puro_R and the Zymo DNA Clean & Concentrator Kit purified PCR product was sequenced. The PCR conditions were: 98 °C for 2 min; 30 cycles of 98 °C for 5 s, 55 °C for 10 s, and 72 °C for 40 s; and, followed by 72 °C for 5 min. Sequencing results confirmed that *520-522* or *602-605* was inserted into the JCVI-syn3.0 genome without mutation. Primers used are listed in Tables S2.

### Construction of plasmid Pmod2-loxpurolox-sp-cre_sp_mCherry

Pmod2-loxpurolox-sp-cre_sp_mCherry was constructed by inserting the spiralen promoter and *mCherry* gene adjacent to the puromycin resistance marker in Pmod2-loxpurolox-sp-cre via Gibson Assembly. The spiralen promoter and *mCherry* gene were amplified using Q5 master mix (New England Biolabs), with the plasmid pTF20mChloxp (gift from Kevin Dybvig) as the template. The linear vector Pmod2-loxpurolox-sp-cre was amplified using Q5 master mix, with plasmid Pmod2-loxpurolox-sp-cre as the template. Primers are listed in Table S2. The expected PCR product sizes listed in Table S3 were confirmed.

### Insertion of mCherry into the JCVI-syn1.0 and RGD6 genomes

The mCherry coding sequence was introduced at random chromosomal locations into JCVI-syn1.0 or RGD6 cells and selected using the puromycin resistance marker, as described previously (Karas et al., 2014). Briefly, the coding sequence of mCherry (Shaner et al., 2004) was introduced into the pLS-Tn5-Puro vector, and a linearized product was then amplified and combined with Tn5 transposase. The resulting transposome was transformed into cells and plated on selective agar medium. Single colonies were picked and grown in liquid culture without selection and screened for suitable mCherry expression using fluorescence microscopy. JCVI-syn1.0+mCherry and RGD6+mCherry clonal isolates with bright fluorescence signals and stable expression of mCherry over many generations were used in the studies reported here.

### Insertion of mCherry into the JCVI-syn3.0 genome

Transformation of JCVI-syn3.0 using Pmod2-loxpurolox-sp-cre_sp_mCherry plasmid was performed as described above. After transforming JCVI-syn3.0 cells with the plasmid Pmod2-loxpurolox-sp-cre_sp_mCherry, diagnostic PCR measurements confirmed delivery of the mCherry and puromycin resistance genes to the genome of JCVI-syn3.0+mCherry. Primers are listed in Table S2.

### Construction of plasmid Pmod2-loxpurolox-sp-cre_538+546-549+592-593+622-623 (clusters 3+4+5+8)

The plasmid Pmod2-loxpurolox-sp-cre_*538*+*546-549*+*592-593*+*622-623* was constructed by inserting the genes *538, 546-549, 592-593*, and *622-623* adjacent to the puromycin resistance marker in Pmod2-loxpurolox-sp-cre via Gibson Assembly. The region containing genes *538, 546-549, 592-593*, and *622-623* was amplified using primers 622 F 6.9K and 592R 7.5 K, PrimeSTAR Max Premix, and plasmid Pmod2-hisarscen-*527*_*538*_*546-549*_*592-593*_*622-623* as the template. The linear vector Pmod2-loxpurolox-sp-cre was PCR amplified using primers vec F 7.5 K and vec R 6.9K, PrimeSTAR Max Premix, and the plasmid Pmod2-loxpurolox-sp-cre as the template. The PCR conditions were: 98 °C 3 min; 30 cycles of 98 °C for 10 s, 55 °C for 10 s, and 72 °C for 1 min; and, followed by 72 °C for 2 min. The PCR amplicon was purified using a Zymo DNA Clean & Concentrator Kit.

Purified DNA fragment *538*_*546-549*_*592-593*_*622-623* and linear vector were assembled via Gibson Assembly. The assembly mix was transformed using competent cell DH5alpha C2987H, grown at 37 °C for 1 h, and plated on an AmpR LB plate. The plate was incubated at 37 °C overnight. Colony PCR was used to screen positive colonies containing the gene *538*_*546-549*_*592-593*_*622-623* insertion in the vector, using Qiagen Multiplex PCR Kit according to the manufacturer’s protocol. The expected sizes of PCR products listed in Table S3 were obtained. Primers for PCR amplification, colony PCR, and sequencing are listed in Table S2. Sanger sequencing data confirmed that the inserted genes in the plasmid had no mutations.

### Construction of JCVI-syn3.0+12345678

Pmod2loxploxcre-*538*+*546*-*549*+*592-593*+*622-623* was used to transform JCVI-syn3.0+1267 (Fig 5C) to obtain JCVI-syn3.0+12345678 (Fig 5C). Junction PCR and Sanger sequencing showed that genes *puro* and *538*+*546*-*549*+*592-593*+*622-623* were inserted in the landing pad of JCVI-syn3.0+1267. The presence of the complete JCVI-syn3.0 genome was confirmed by colony multiplex PCR. Genomic DNA was extracted from the strain JCVI-syn3.0+12345678 and used as the template for PCR amplification of each gene. PrimeSTAR Max Premix, primers 7.8k LJF(401) and 520-602jR(326), 520-602jF(326) and 7.8k ploAR, 7.5k LJF(190) and 7.5k2jR(517), and seq_F1 and puro_R, respectively, were used to amplify regions containing each gene. PCR products were purified using a Zymo DNA Clean & Concentrator Kit. Sanger sequencing of PCR products, using primers for 7.5k in Table S2, excluding 7.5 K LJ F(190), revealed a single point mutation (G to T) in the intergenic region 5’ of gene *546*.

### Construction of plasmid Pmod2-loxpurolox-sp-cre_546-549+622-623 (clusters 4+8)

The plasmid Pmod2-loxpurolox-sp-cre_*546-549*+*622-623* was constructed by inserting the genes *546-549*, and *622-623* adjacent to the puromycin resistance marker in Pmod2-loxpurolox-sp-cre via Gibson Assembly. The genes *546-549* were amplified using Q5 master mix, with JCVI-syn1.0 genome as the template, as well as primers 546 F to 622 and 546 R to 622. The genes *622-623* were amplified using Q5 master mix, with JCVI-syn1.0 as the template, and primers 622 F 6.9K and 622 R to 546. The linear vector Pmod2-loxpurolox-sp-cre was amplified using the plasmid Pmod2-loxpurolox-sp-cre as the template, Q5 master mix, and primers vec F 546-622 and vec R 6.9K. The PCR conditions were: 98 °C 3 min; 30 cycles of 98 °C for 10 s, 58 °C for 15 s, and 72 °C for 2.5 min; and, followed by 72 °C for 3 min.

DNA fragments *546-549, 622-623* and linear vector were assembled via Gibson Assembly. The assembly mix was transformed using competent cell DH5alpha C2987H, grown at 37 °C for 1 h and plated on an AmpR LB plate. The plate was incubated at 37 °C overnight. Colony PCR was used to screen positive colonies with a gene *546-549_622-623* insertion in the vector using a Qiagen Multiplex PCR Kit protocol. The expected DNA sizes were listed in Table S3. Primers for PCR amplification, colony PCR and sequencing are listed in Table S2. Sanger sequencing confirmed that the plasmid has no mutations.

### Construction of strain JCVI-syn3.0+124678

Pmod2loxploxcre-*546-549*+*622-623* was used to transform JCVI-syn3.0+1267 to obtain JCVI-syn3.0+124678 (Fig 5C). Junction PCR and Sanger sequencing showed that gene puro and *546*-*549*+*622-623* were inserted in the landing pad of JCVI-syn3.0+1267. These strains were confirmed to have the complete JCVI-syn3.0 genome by colony multiplex PCR.

Genomic DNA was extracted from strain JCVI-syn3.0+124678. PrimeSTAR Max Premix, as well as primers seq_F1 and puro_R, were used to amplify added genes using the genomic DNA of JCVI-syn3.0+124678 as the template. PCR products were purified using a Zymo DNA Clean & Concentrator Kit. Sanger sequencing, using sequencing primers in Table S2, confirmed the absence of any mutations.

### Editing mycoplasmal chromosomes in yeast by CRISPR/Cas9 and genome transplantation

To put back several gene clusters in the genome of JCVI-syn3.0 in different locations, we used CRISPR/Cas9 to edit the genome of the mycoplasma in yeast. Editing the genome as a yeast centromeric plasmid clone (YCp) is required to delete genes from any synthetic strain or to insert genes at sites other than the dual *loxP* landing pad. After editing the genome as a YCp, the edited genome is transplanted into an *M. capricolum* recipient strain to render a mycoplasmal transplant programmed solely by the new chromosome.

The *S. cerevisiae* yeast strain VL6_48N_cas9 was obtained from Dr. Daniel G. Gibson (Codex DNA, Inc.). VL6_48N_cas9 carries the cas*9* gene in a yeast chromosome, which constitutively expresses Cas9 as described previously (Kannan et al., 2016). Yeast strains were grown in the yeast media “BD Difco™ YPD Broth (Fisher Scientific) plus 60 mg of adenine per liter,” “c7112, CM Glucose Broth, Dry, Adenine-60, without Histidine,” “c0230, CM Glucose Agar, Dry, w/o Histidine, Tryptophan,” or “c7221, CM Glucose media, Dry, w/o Histidine, Uracil, 2 % (w/v) agar added to make agar plate.” These media were all obtained from Teknova, unless indicated otherwise.

### Insertion of JCVI-syn3.0 and JCVI-syn3A bacterial genomes into yeast

To insert bacterial genomes into yeast, fusion of JCVI-syn3.0 or JCVI-syn3A with yeast strain VL6_48N_cas9 was performed as described elsewhere (Karas et al., 2013). Briefly, yeast VL6_48N_cas9 was grown overnight in YPD growth medium supplemented with adenine at 120 mg/L until OD600 ≈(1.0 to 2.0). Cells were centrifuged, and the supernatant was discarded. Cells were then washed in water and 1 mol/L sorbitol, followed by resuspension and centrifugation. The supernatant was discarded, and the cell pellet was then resuspended in SPEM solution, consisting of 1 mol/L sorbitol, 10 mmol/L EDTA, 2.08 g/L Na2HPO4·7H2O, 0.32 g/L NaH2PO4·1H2O, 30 µL of β-mercaptoethanol (Sigma-Aldrich), and 40 µL of Zymolyase-20T solution (200 mg Zymolyase (USB), 1 mL of 1 mol/L Tris-HCl pH 7.5, 10 mL 50 % (v/v) glycerol, and 9 mL dH2O). After a 40 min incubation, the resulting yeast spheroplasts were ready for fusion to bacteria.

A 5 mL culture of JCVI-syn3.0 or JCVI-syn3A was grown overnight to pH (6.7 to 7.0). Chloramphenicol was then added to a final concentration of 100 mg/L, and cells were incubated at 37 °C for 1.5 h. Cells were subsequently centrifuged and resuspended in 50 µL of 0.5X resuspension buffer, where 1X resuspension buffer is composed of 0.5 mol/L sucrose, 10 mmol/L Tris HCl pH 7.5, 10 mmol/L CaCl2, and 2.5 mmol/L MgCl2, with pH adjusted to pH 7.5.

200 µL of yeast spheroplasts and 50 µL of chloramphenicol-treated JCVI-syn3.0 or JCVI-syn3A cells were mixed gently. 1 mL of 20 % (w/v) PEG 8000 (USB) solution, consisting of 20 % (w/v) PEG 8000, 10 mmol/L Tris-HCl pH 8.0, 10 mmol/L CaCl2, and 2.5 mmol/L MgCl2, with pH adjusted to pH 8.0, was added to the yeast and mycoplasma mixture. After gentle mixing and further incubation at room temperature for 20 min, cells were centrifuged at 1500 g for 7 min and the supernatant removed. The cell pellet was resuspended in 1 mL of SOS, composed of 1 mol/L sorbitol, 6 mmol/L CaCl2, 2.5 g/L yeast extract, and 5 g/L Bacto Peptone, and incubated for 30 min at 30 °C. The 1 mL cell suspension in SOS was mixed with top agar and plated on agar plates containing CM Glucose Broth, Dry, Adenine-60, without Histidine with 1 mol/L sorbitol. Yeast transformants appeared after (3 to 5) days.

### Yeast assembly of Pmod2-hisarscen-610_520-522_602-605 (clusters 1+6+7)

Adding single gene clusters to JCVI-syn3.0 did not rescue the pleomorphic phenotype of JCVI-syn3.0. We therefore added multiple gene clusters to JCVI-syn3.0 in an attempt to correct this phenotype. We assembled several gene clusters in yeast. The genes *610, 520-522, 602-605* were amplified using amplification primers listed in Table S2 and PrimeSTAR Max Premix, with JCVI-syn1.0 genome as the template. The vector Pmod2-hisarscen was amplified using PrimeSTAR Max Premix with plasmid Pmod2-hisarscen-cm as the template, as well as primers Y 610 vec F and Y vec R in Table S2. The PCR conditions were: 98 °C for 2 min; 30 cycles of 98 °C for (5 or 10) s, 55 °C for 10 s, and 72 °C for (40 or 60) s; and, followed by 72 °C for 3 min. The plasmid Pmod2-hisarscen-cm was constructed by assembly of *his, ars, cen*, and *cm* genes with vector Pmod2-MCS (Epicentre) via Gibson Assembly. The primers used for construction of Pmod2-hisarscen-cm are listed in Table S2. (25 to 50) ng of Zymo DNA Clean & Concentrator Kit purified DNA fragment *610, 520-522, 602-605* and linear vector Pmod2-hisarscen, with 50 bp overlapping with adjacent fragments, were assembled in yeast strain VL6_48 using the lithium acetate PEG method (Gietz and Schiestl, 2007). Plasmid was isolated from yeast, then transformed and propagated in *E. coli* STbl4 (Life Technology). Plasmid was then extracted for Sanger sequencing, which subsequently confirmed the absence of mutations. PCR amplification primers, junction primers for diagnostic PCR, and sequencing primers are listed in Tables S2.

### Yeast assembly of Pmod2-hisarscen-527_538_546-549_592-593_622-623 (clusters 2+3+4+5+8)

The gene clusters *527, 538, 546-549, 592-593* and *622-623* were amplified and assembled in yeast as described in the previous section. Diagnostic PCR with junction primers gave products with the expected sized indicated in Table S3, thereby confirming the correct assembly of genes *527, 538, 546-549, 592-593, 622-623*. Propagation of the plasmid in *E. coli* STbl4, followed by Sanger sequencing, confirmed the absence of mutations. PCR amplification primers, junction primers for diagnostic PCR, and sequencing primers are listed in Table S2.

### CRISPR/Cas9 guide RNA production and quantification

We used CRISPR/Cas9 methods to insert genes into the JCVI-syn3.0 genome residing in the yeast strain VL6_48N_cas9_ JCVI-syn3.0. Initially, we designed single guide RNA (sgRNA) targeting the gene(s) of interest in the JCVI-syn3.0 genome. A (119 to 120) bp dsDNA was then obtained via a PCR reaction containing the following: 1 µL of the guide RNA forward primer 7.8k_gRNA_F (10 µmol/L) or 7.5k_gRNA_F (10 µmol/L), which included the T7 promoter, 19-20 bp guide RNA target, and an overlap with 83 bp of sgRNA_Template (Mali et al., 2013), 1 µL of reverse primer gRNA_R (10 µmol/L), 10 ng of 83 bp sgRNA template, and Q5 master mix or 2X PrimeSTAR Max Premix (Takara) in a 20 µL PCR reaction. The PCR conditions were: 2 min at 98 °C; 30 cycles of 98 °C for 10 s, 55 °C for 10 s, and 72 °C for 10 s; and, followed by 72 °C for 5 min. We then purified the PCR amplicons using a Zymo DNA Clean & Concentrator Kit. We used 8 µL of the PCR amplicons as template in a 20 µL transcription reaction with the T7 RiboMAX™ Express Large Scale RNA Production System according to the manufacturer’s instructions. Briefly, we combined 10 µL of RiboMAX™ Express T7 2X Buffer, 158 ng of the previously generated PCR amplicon (in 8 µL H2O), and 2 µL Enzyme Mix T7 Express. The transcription reaction was incubated at 37 °C for 4 hr or overnight, followed by addition of 1 µL of Turbo DNase (Thermo Fisher Scientific) provided with the kit and incubation at 37 °C for another 15 min. Following a standard protocol, the reaction volume was adjusted to 750 µL with RNase-free water and 0.1 volumes of 3M sodium acetate was added, followed by phenol-chloroform extraction. RNA was then precipitated with two volumes 100 % (v/v) ethanol and washed with 500 µL 70 % (v/v) ethanol. The pellet was air-dried and then resuspended in 40 µL RNase-free water. The Qubit™ RNA HS Assay Kit (Thermo Fisher Scientific) was used to quantify guide RNA. Primers used are listed in Table S2.

### CRISPR donor DNA preparation for yeast strain VL6_48N_cas9_JCVI-syn3.0+1267

Gene *527* was inserted into VL6_48N_cas9_JCVI-syn3.0 between gene *526* and adjacent gene *528* via CRISPR/Cas9. Gene *527* was amplified using PrimeSTAR Max Premix and primers 527_50bp_F and 7.5k_ 50bp_R with 50 bp overlapping each side of insertion sites on the JCVI-syn3.0 genome. Genes *610, 520-522, 602-605* were inserted in VL6_48N_cas9_ JCVI-syn3.0 between genes *0609* and adjacent gene *611* via CRISPR/Cas9. Genes *610, 520-522, 602-605* were amplified using PrimeSTAR Max Premix with Pmod2-hisarscen-*610—520-522—602-605* as the template, and primers 610_7.8k_F and 610_7.8k_R with 50 bp overlapping each side of insertion sites on the JCVI-syn3.0 genome. The presence and location of genes were confirmed by obtaining the expected PCR products across junctions (Table S3). Primers are listed in Table S2.

### CRISPR donor DNA preparation for yeast strain VL6_48N_cas9_JCVI-syn3.0+2

Gene *527* was inserted into VL6_48N_cas9_JCVI-syn3.0 between gene *0526* and adjacent gene *0528* via CRISPR/Cas9. Gene *0527* was amplified using PrimeSTAR Max Premix, primers 7.5 K 2J R(517) and 7.5 K LJ F(190), and Pmod2-hisarscen-*527*_*538*_*546-549*_*592-593*_*622-623* as the template. The expected sizes of PCR products listed in Table S3 were obtained. Primers are listed in Table S2.

### CRISPR donor DNA preparation for yeast strain VL6_48N_cas9_JCVI-syn3.0+234578

Gene *610* was amplified, with plasmid Pmod2-hisarscen-*610—520-522—602-605* as the template, using PrimeSTAR Max Premix and primers 610_7.8k_F and 610_7.8k_R with 50 bp overlapping each side of insertion sites on the JCVI-syn3.0 genome. Gene *610* was inserted in VL6_48N_cas9_ JCVI-syn3.0 after gene *609* and before gene *611* via CRISPR/Cas9. The genes *527, 538, 546-549, 592-593*, and *622-623* were inserted into VL6_48N_cas9_JCVI-syn3.0 after gene *526* and before *528* via CRISPR/Cas9. Genes *527, 538, 546-549, 592-593*, and *622-623* were amplified with plasmid Pmod2-hisarscen-*527_538_546-549_592-593_622-623* as the template, PrimeSTAR Max Premix, and primers 527_50bp_F and 7.5k_ 50bp_R with 50 bp overlapping each side of insertion sites on the JCVI-syn3.0 genome. Primers are listed in Table S2.

For CRISPR donor DNA preparation, the PCR conditions were: 2 min at 98 °C; 30 cycles of 98 °C for 10 s, 55 °C for 10 s, and 72°C for 1 min; and, followed by 72 °C for 3 min.

### Transformation and screening positive yeast transformants for gene insertions

The transformation of yeast, selection, and screening for positive colonies were performed as described elsewhere (Kannan et al., 2016) with some changes. Briefly, 500 ng of donor DNA, (400 to 500) ng guide RNA, and 100 ng PCC1BAC_trp plasmid (GenBank accession MN982904) as the selective trp marker (gift from Dr. Billyana Tsvetanova, Synthetic Genomics, Inc.) were co-transformed with yeast VL6_48N_cas9_JCVI-syn3.0. After electroporation, 1 mL of YPDA/sorbitol media was mixed with cells and transferred to a 30 °C shaker for 2 h. 100 µL of culture was plated on selection plates containing CM Glucose Agar, Dry, w/o Histidine, Tryptophan. Colonies appeared after 4 days and diagnostic PCR was run with junction primers to screen positive colonies with genomic modification. The expected sizes of PCR products listed in Table S3 were obtained. For positive clones, multiplex PCR was run to confirm genome integrity. Primers for junction and multiplex PCR are listed in Table S2.

### Genome transplantation

Mycoplasmal genomes that are maintained and manipulated in yeast as yeast centromeric plasmids can be booted up by genome transplantation into a recipient organism, *Mycoplasma capricolum*, as described in detail elsewhere (Gibson et al., 2010; Hutchison et al., 2016; Lartigue et al., 2009; Tsarmpopoulos et al., 2016). The positive yeast clones containing verified chromosomes were transplanted, as described previously (Lartigue et al., 2009). Single colonies isolated from mycoplasmal transplant outgrowth populations (grown under selection on tetracycline plates) were transferred to SP4 liquid culture without tetracycline. After growth for 3 days, cells were imaged as wet mounts using DIC microscopy to determine morphology.

To confirm the chromosomal constructs in mycoplasmal transplants, genomic DNA was extracted from cells in mid-log phase liquid culture and used as PCR template. The inserted genes were amplified using PCR with PrimeSTAR Max Premix. The PCR conditions were: 2 min at 98 °C; 30 cycles of 98 °C for 10 s, 55 °C for 10 s, and 72 °C for 1 min; and, followed by 72 °C for 3 min. Zymo DNA Clean & Concentrator Kit purified PCR products were checked using Sanger sequencing and no mutation was found in any of the constructs. The sequencing primers used for these transplants are listed in Table S2.

To confirm genome integrity, Qiagen Multiplex PCR Kit was used for colony PCR, using primers listed in Table S2. Each pair of forward and reverse primers generated a PCR product specific for a genome segment. The expected product lengths are indicated. The PCR conditions were: 95 °C for 15 min; 34 cycles of 94 °C for 30 s, 52 °C for 90 s, and 68 °C for 2 min; followed by 68 °C for 3 min. For all PCR, the primers were diluted to 2-10 µmol/L and the final concentration of each primer is 0.1-0.5 µmol/L. Genome manipulation in yeast and transplantation do not induce mutations as shown by sequencing before and after transplantation (Lartigue et al., 2009), and cells were passaged as few generations as possible – typically ≈20 generations – before assaying cell morphology. For experiments using CRISPR/Cas9, guide RNA forward primers and donor DNA oligonucleotides were ordered from IDT.

### Gene removal from the JCVI-syn3A genome in yeast

CRISPR/Cas9 methods were used to delete the following individual gene clusters from VL6_48N_cas9_JCVI-syn3A: *520-522, 610, 602-605, 527, 538, 546-549, 592-593*, or *622-623*. For the methods of production of guide RNA targeting each gene cluster see *CRISPR/Cas9 guide RNA production and quantification* above.

Single stranded donor DNA 100 base oligonucleotides (IDT) were used as a patch to seal the gap after deleting genes *520-522*. The 100 base donor carried two 50 bp overlaps with the two sides of genomic DNA breaks. PCC1BAC_trp plasmid (100 ng), 1 µg of donor DNA, and 500 ng of guide RNA targeting gene *520-522* (see *CRISPR/Cas9 guide RNA production and quantification*) were co-transformed into yeast competent cell VL6_48N_cas9_syn3A. Later positive colonies were screened using a Qiagen Multiplex PCR Kit. The expected DNA sizes are listed in Table S3. Gene *522* and its 5’ flanking region, gene *521*, and most of *520* were deleted and we obtained VL6_48N_cas9_syn3AΔ*520-522*.

Using the same methods, we obtained the following strains:

- VL6_48N_cas9_syn3AΔ*527* with gene *527* deleted from JCVI-syn3A
- VL6_48N_cas9_syn3AΔ*538* with gene *538* deleted from JCVI-syn3A
- VL6_48N_cas9_syn3AΔ*546-549* with gene *546-549* deleted from JCVI-syn3A
- VL6_48N_cas9_syn3AΔ*592-593* with gene *592-593* deleted from JCVI-syn3A
- VL6_48N_cas9_syn3AΔ*602-605* with gene *602-605* deleted from JCVI-syn3A
- VL6_48N_cas9_syn3AΔ*610* with gene *610* deleted from JCVI-syn3A
- VL6_48N_cas9_syn3AΔ*622-623* with gene *622-623* deleted from JCVI-syn3A

The expected DNA sizes were listed in Table S3. For positive clones with the desired gene deletion, multiplex PCR using a Qiagen Multiplex PCR Kit was performed to confirm the whole genome integrity, using primers listed in Table S2.

Genome transplantation was carried out to recover the modified JCVI-syn3A strains from yeast and diagnostic PCR and multiplex PCR were used to confirm the insert junctions and genome integrity. Junction primers are listed in Table S2. Primers for gene removal and primers for JCVI-syn3A genome integrity used are listed in Table S2, and the expected DNA amplicons listed in Table S3 were obtained.

### Deletion of one gene cluster from JCVI-syn3.0+1267

Strain JCVI-syn3.0+*527*+*610*+*520-522*+*602-605* showed the JCVI-syn3A phenotype. We tried to delete one gene cluster from the strain to obtain the minimal gene number needed to restore the JCVI-syn3A phenotype.

We used CRISPR/Cas9 methods to delete genes from yeast strain VL6_48N_Cas9_JCVI-syn3.0+*527*+*610*+*520-522*+*602-605*. We obtained guide RNA first. To make 520 gRNA2, we designed single guide RNA (sgRNA) targeting the gene *520-522* in JCVI-syn3.0 genome. (119 to 120) bp dsDNA was obtained via PCR with the guide RNA forward primer 520 gRNA2_F (with T7 promoter, (19 to 20) bp guide RNA target, and overlap with 83 bp sgRNA template (Mali et al., 2013)), reverse primer gRNA_R, 83 bp sgRNA template, and PrimeSTAR Max Premix in a 20 µL PCR reaction. The (119 to 120) bp dsDNA was transcribed *in vitro* using T7 RiboMAX™ Express Large Scale RNA Production System (Promega Corporation) in 20 µL volumes, including 10 µL 2X buffer, 2 µL of T7 RNA polymerase, and 200 ng dsDNA. Guide RNA was purified using Acid-Phenol:Chloroform at pH 4.5 (Thermo Fisher Scientific), precipitated using 100 % (v/v) ethanol and 3M sodium acetate, and dissolved in 40 µl RNase free water. A Qubit™ RNA HS Assay Kit (Thermo Fisher Scientific) was used to quantify guide RNA.

To make VL6_48N_Cas9_JCVI-syn3.0+*527*+*520-522*+*602-605*+*610*Δ*520-522* (Table S4), gene cluster *520-522* was deleted from yeast strain VL6_48N_Cas9_JCVI-syn3.0+*527*+*520-522*+*602-605*+*610* using CRISPR/Cas9 methods. Briefly. 520 gRNA2 and 520gRNA3 were used to guide CAS9 protein to the gene *520-522* target sites and cut the genome of JCVI-syn3.0+*527*+*520-522*+*602-605*+*610*, 520 patch was used as the donor DNA to seal the cut, and PRS316 plasmid (bearing the ura3 marker) was used as the selective marker to co-transform the competent yeast strain VL6_48N_cas9_JCVI-syn3.0+*527*+*520-522*+*602-605*+*610*. The transformation reaction mix was incubated at 30 °C for 2 h and plated for selection on CM Glucose Broth without Histidine and Uracil c7221 agar medium. The plate was incubated at 30 °C for (3 to 4) days, and diagnostic PCR using junction primers was run to screen for positive colonies that had the gene cluster *520-522* deleted using a Qiagen Multiplex PCR Kit. PCR products with the expected sizes listed in Table S3 were obtained.

Using the same methods, we deleted gene clusters *527, 602-605* or *610*, to obtain strains JCVI-syn3.0+*527*+*520-522*+*602-605*+*610*Δ*527*, JCVI-syn3.0+*527*+*520-522*+*602-605*+*610*Δ*602-605*, and JCVI-syn3.0+*527*+*520-522*+*602-605*+*610*Δ*610* (Table S4), respectively. The guide RNA forward primers, donor DNA, primers for JCVI-syn3.0 genome integrity, and junction primers are listed in Table S2. PCR products of the expected sizes listed in Table S3 were obtained.

### Deletion of genes from JCVI-syn3.0+126 using CRISPR/Cas9

Strain JCVI-syn3.0+*527*+*520-522*+*602-605* showed a JCVI-syn3A phenotype. Deletion of gene *527* from JCVI-syn3.0+*527*+*610*+*520-522*+*602-605* caused a reversion to a pleomorphic phenotype, thereby indicating that gene *527* is necessary to confer the JCVI-syn3A phenotype in that strain. Similarly, deletion of the individual gene clusters *520-522* or *602-605* from the strain also rendered the pleomorphic cell phenotype. We therefore sought to determine the effect of removing the individual genes within these three clusters on the JCVI-syn3A phenotype of strain JCVI-syn3.0+*527*+*520-522*+*602-605*. CRISPR/Cas9 methods were used in a yeast strain carrying JCVI-syn3.0+*527*+*520-522*+*602-605* to delete single genes, or gene pairs, from the genome of JCVI-syn3.0+*527*+*520-522*+*602-605* in yeast. The yeast strain VL6_48N_cas9_JCVI-syn3.0+*527*+*520-522*+*602-605* (carrying the *trp* plasmid and *ura3* plasmid from previous experiments) was passaged twice (with 1:1000 dilution), grown in CM Glucose Broth without Histidine liquid media for 24 h, and then diluted and plated on separate agar plates containing CM Glucose Broth: (i) without Histidine; (ii) without Histidine and Tryptophan; or, iii) without Histidine and Uracil. Yeast colonies growing only on CM Glucose Broth without Histidine were picked. Trp and ura3 markers were absent from these clones, and one was used for the following experiments.

For gene cluster *520-522*, there were 6 deletion combinations, including deletion of gene *520, 521* or *522, 520*+*521, 521*+*522*, or *520*+*522*. For gene cluster *602-605*, there are also 6 deletion combinations, including deletion of gene *602, 604, 605, 602-604, 604+605*, or *602*+*605*. Note that locus tag _0603 is no longer annotated as a gene; rather, it represented a small region of DNA 5’ of gene *604*.

To generate some mutants, we used the same CRISPR/Cas9 methods to remove genes, except that donor DNA was produced via PCR. To make JCVI-syn3.0+*527*+*520-522*+*602-605*Δ*521*, we used primers 521-40bpR and 520 jump F1, JCVI-syn1.0 genomic DNA as the template, and PrimeSTAR Max Premix in a PCR to produce a 730 bp DNA donor. To make JCVI-syn3.0+*527*+*520-522*+*602-605*Δ*602*Δ*605* or JCVI-syn3.0+*527*+*520-522*+*602-605*Δ*605*, we used PCR primers 604-40-F1 and 604-40-R1, JCVI-syn1.0 genomic DNA as template, and PrimeSTAR Max Premix. To make JCVI-syn3.0+*527*+*520-522*+*602-605*Δ*520*Δ*522*, we used PCR primers 521-40bp-R, 521-40bp-F, JCVI-syn1.0 genomic DNA as the template, and PrimeSTAR Max Premix to produce a 760 bp DNA donor. To make JCVI-syn3.0+*527*+*520-522*+*602-605*Δ*604*, a 500 bp donor DNA was made via a fusion PCR. First, we used primers 602jumpF2(315) and 604-40bpR, to obtain a 516 bp PCR product. Next, the PCR product was purified using a Zymo DNA Clean & Concentrator Kit. At the same time, primers 604-80bp and 602jumpR3(492), Syn1.0 genomic DNA as the template, and PrimeSTAR Max Premix were used for PCR to yield a 480 bp PCR product, which was the purified using a Zymo DNA Clean & Concentrator Kit. 20 ng of each of the above two PCR products, primers 602jumpF2(315) and 602jumpR3(492), and PrimeSTAR Max Premix were used to produce 996 bp donor DNA. The PCR conditions were: 98 °C for 3 min; 30 cycles of 98 °C for 10 s, 55 °C for 10 s, 72 °C for 1 min, and, followed by 72 °C for 2 min. The PCR amplicon was purified using a Zymo DNA Clean & Concentrator Kit. 500 ng each of guide RNA 602gRNA3, 605gRNA1, 604gRNA2, as well as 1 µg of donor DNA and 100 ng PCC1BAC_trp plasmid, were used to transform yeast competent cells.

To delete gene *520* from the genome of JCVI-syn3.0+*527*+*520-522*+*602-605* in yeast (Table S4), 520gRNA3 was used to guide CAS9 protein to the 520gRNA3 target site and cut the genome of JCVI-syn3.0+*527*+*520-522*+*602-605*. Single-stranded oligonucleotide 520-80 base patch was used as donor DNA to seal the cut, and PCC1BAC_trp plasmid was used as the selective marker to co-transform the competent yeast strain VL6_48N_cas9_JCVI-syn3.0+*527*+*520-522*+*602-605*. The transformation reaction mix was incubated at 30 °C for 2 h and plated on a selective plate with CM Glucose Broth without Histidine and Tryptophan. The plate was incubated at 30 °C for (3 to 4) days, and diagnostic PCR using junction primers was run to screen positive colonies that had the gene *520* deleted. The expected DNA sizes of PCR products listed in Table S3 were obtained. The yeast strain with gene *520* deleted was used for genome transplantation to generate a mycoplasmal transplant with the genome of JCVI-syn3.0+*527*+*520-522*+*602-605*Δ*520*. Junction PCR results confirmed that gene *520* was deleted.

Using the same method, we obtained the following strains:

- JCVI-syn3.0+*527*+*520-522*+*602-605*Δ*602*
- JCVI-syn3.0+*527*+*520-522*+*602-605*Δ*602-604*
- JCVI-syn3.0+*527*+*520-522*+*602-605*Δ*604-605*
- JCVI-syn3.0+*527*+*520-522*+*602-605*Δ*522*
- JCVI-syn3.0+*527*+*520-522*+*602-605*Δ*520-521*
- JCVI-syn3.0+*527*+*520-522*+*602-605*Δ*521-522*

**Table S2. (Excel spreadsheet)** lists primers for JCVI-syn3.0 genome integrity analysis and junction primers.

**Table S3.** lists expected DNA sizes for diagnostic PCR screening of gene deletion.

**Table S3.**
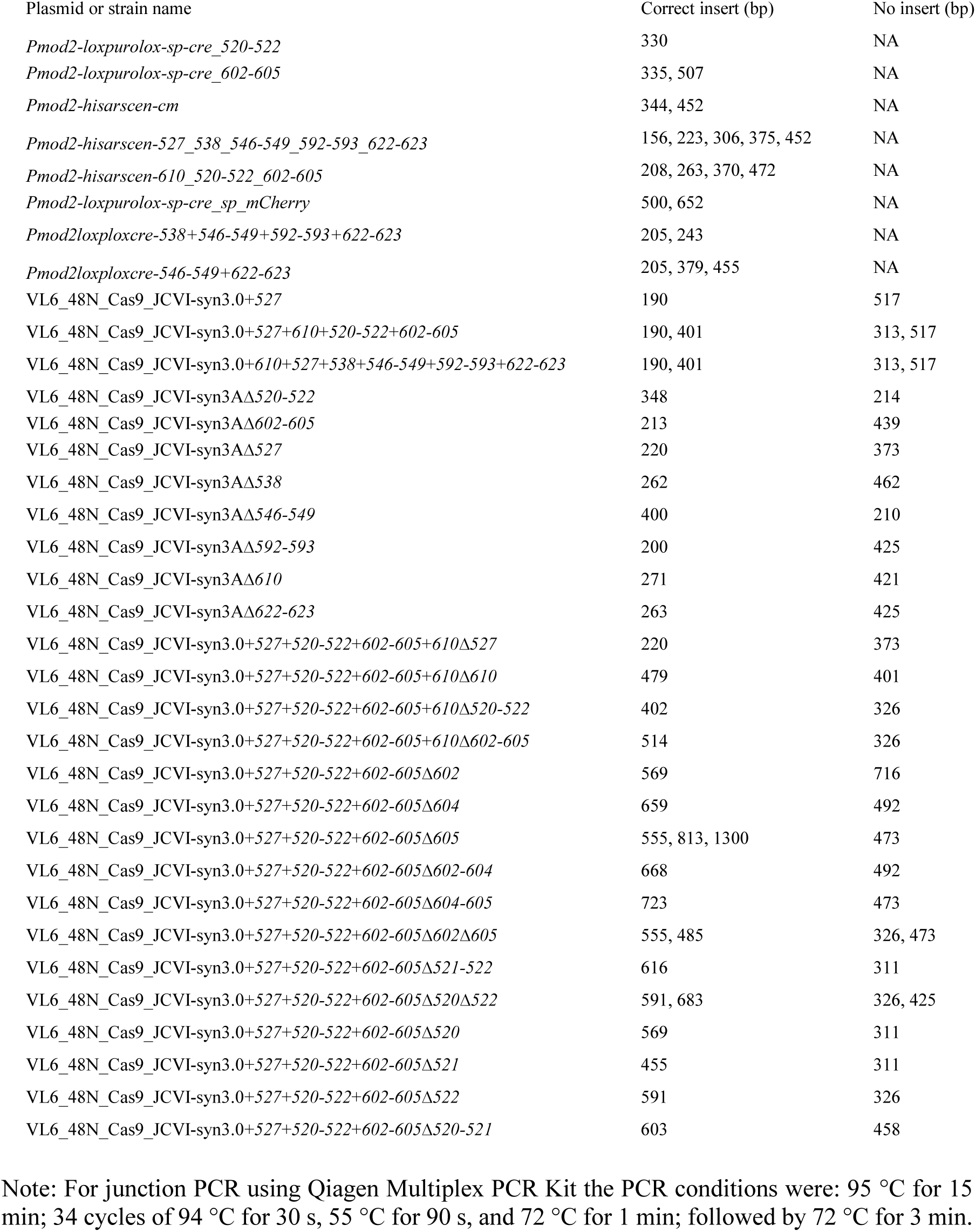
Expected sizes of amplicons using junction primers for strains.

**Table S4.**
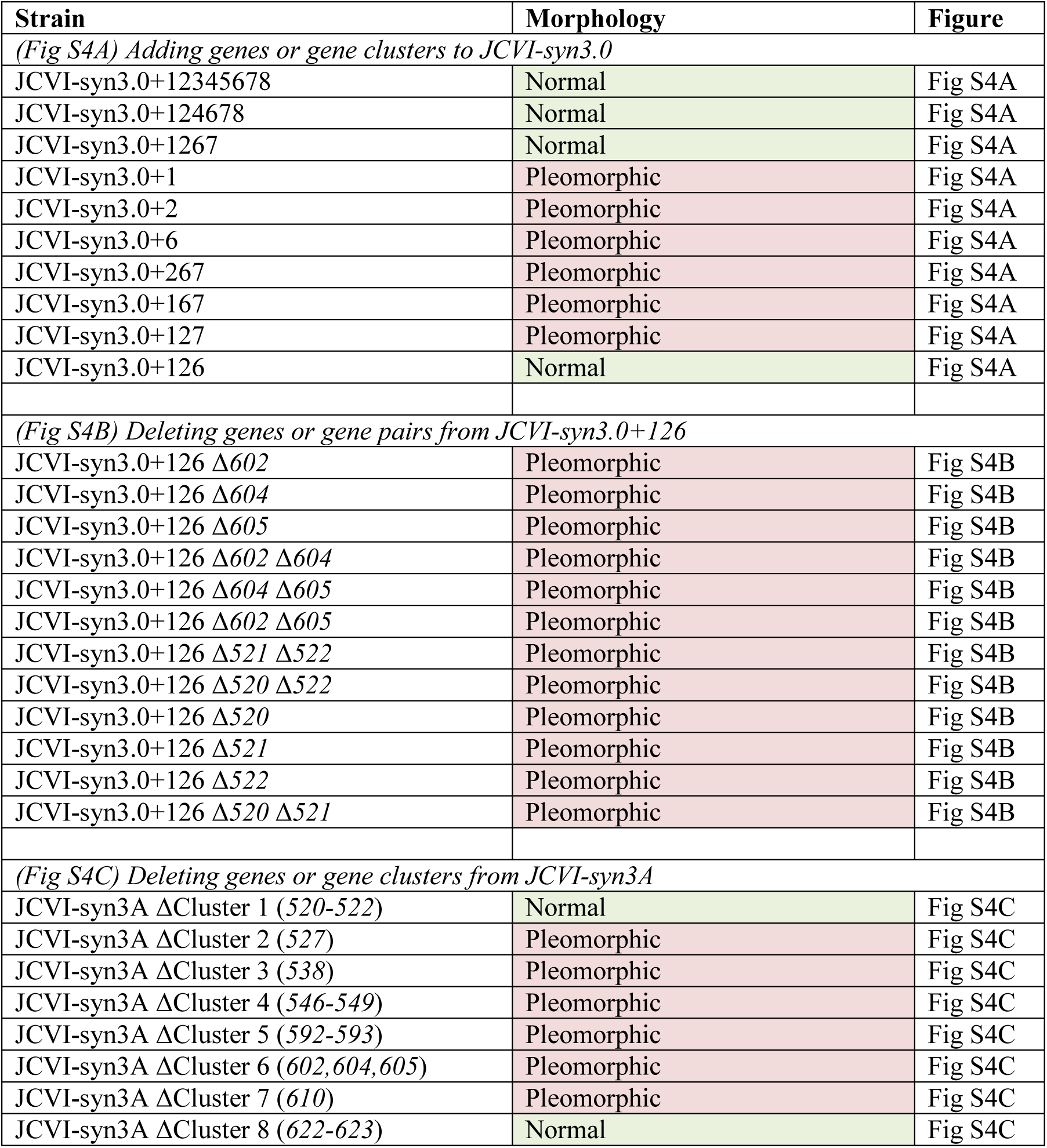
Strains constructed to determine genes required for normal morphology. (A) Additions of genes or gene clusters to JCVI-syn3.0. These strains are shown in Fig S4A. Addition of cluster 1 (*520-522*), cluster 2 (*527*), and cluster 6 (*602,604,605*) was necessary to restore a nearly normal morphology in JCVI-syn3.0+126. These are the same strains listed in Fig 5C of the main text. (B) Deletion of genes or gene pairs from JCVI-syn3.0+126. These strains are shown in Fig S4B. Deletion of any gene caused loss of the nearly normal morphology. (C) Deletion of genes or gene clusters from JCVI-syn3A. These strains are shown in Fig S4C. Surprisingly, JCVI-syn3A retained a nearly normal morphology following deletion of cluster 1 (*520-522*), although cluster 1 was necessary for the nearly normal morphology in JCVI-syn3.0+126.

| Strain | Morphology | Figure |
| --- | --- | --- |
| *(Fig S4A) Adding genes or gene clusters to JCVI-syn3.0* |  |  |
| JCVI-syn3.0+12345678 | Normal | Fig S4A |
| JCVI-syn3.0+124678 | Normal | Fig S4A |
| JCVI-syn3.0+1267 | Normal | Fig S4A |
| JCVI-syn3.0+1 | Pleomorphic | Fig S4A |
| JCVI-syn3.0+2 | Pleomorphic | Fig S4A |
| JCVI-syn3.0+6 | Pleomorphic | Fig S4A |
| JCVI-syn3.0+267 | Pleomorphic | Fig S4A |
| JCVI-syn3.0+167 | Pleomorphic | Fig S4A |
| JCVI-syn3.0+127 | Pleomorphic | Fig S4A |
| JCVI-syn3.0+126 | Normal | Fig S4A |
| *(Fig S4B) Deleting genes or gene pairs from JCVI-syn3.0+126* |  |  |
| JCVI-syn3.0+126 Δ602 | Pleomorphic | Fig S4B |
| JCVI-syn3.0+126 Δ604 | Pleomorphic | Fig S4B |
| JCVI-syn3.0+126 Δ605 | Pleomorphic | Fig S4B |
| JCVI-syn3.0+126 Δ602 Δ604 | Pleomorphic | Fig S4B |
| JCVI-syn3.0+126 Δ604 Δ605 | Pleomorphic | Fig S4B |
| JCVI-syn3.0+126 Δ602 Δ605 | Pleomorphic | Fig S4B |
| JCVI-syn3.0+126 Δ521 Δ522 | Pleomorphic | Fig S4B |
| JCVI-syn3.0+126 Δ520 Δ522 | Pleomorphic | Fig S4B |
| JCVI-syn3.0+126 Δ520 | Pleomorphic | Fig S4B |
| JCVI-syn3.0+126 Δ521 | Pleomorphic | Fig S4B |
| JCVI-syn3.0+126 Δ522 | Pleomorphic | Fig S4B |
| JCVI-syn3.0+126 Δ520 Δ521 | Pleomorphic | Fig S4B |
| *(Fig S4C) Deleting genes or gene clusters from JCVI-syn3A* |  |  |
| JCVI-syn3A ΔCluster 1 (520-522) | Normal | Fig S4C |
| JCVI-syn3A ΔCluster 2 (527) | Pleomorphic | Fig S4C |
| JCVI-syn3A ΔCluster 3 (538) | Pleomorphic | Fig S4C |
| JCVI-syn3A ΔCluster 4 (546-549) | Pleomorphic | Fig S4C |
| JCVI-syn3A ΔCluster 5 (592-593) | Pleomorphic | Fig S4C |
| JCVI-syn3A ΔCluster 6 (602,604,605) | Pleomorphic | Fig S4C |
| JCVI-syn3A ΔCluster 7 (610) | Pleomorphic | Fig S4C |
| JCVI-syn3A ΔCluster 8 (622-623) | Normal | Fig S4C |

**Figure S4. Images of strains to determine genes required for normal morphology.**

**Figure S4A.**
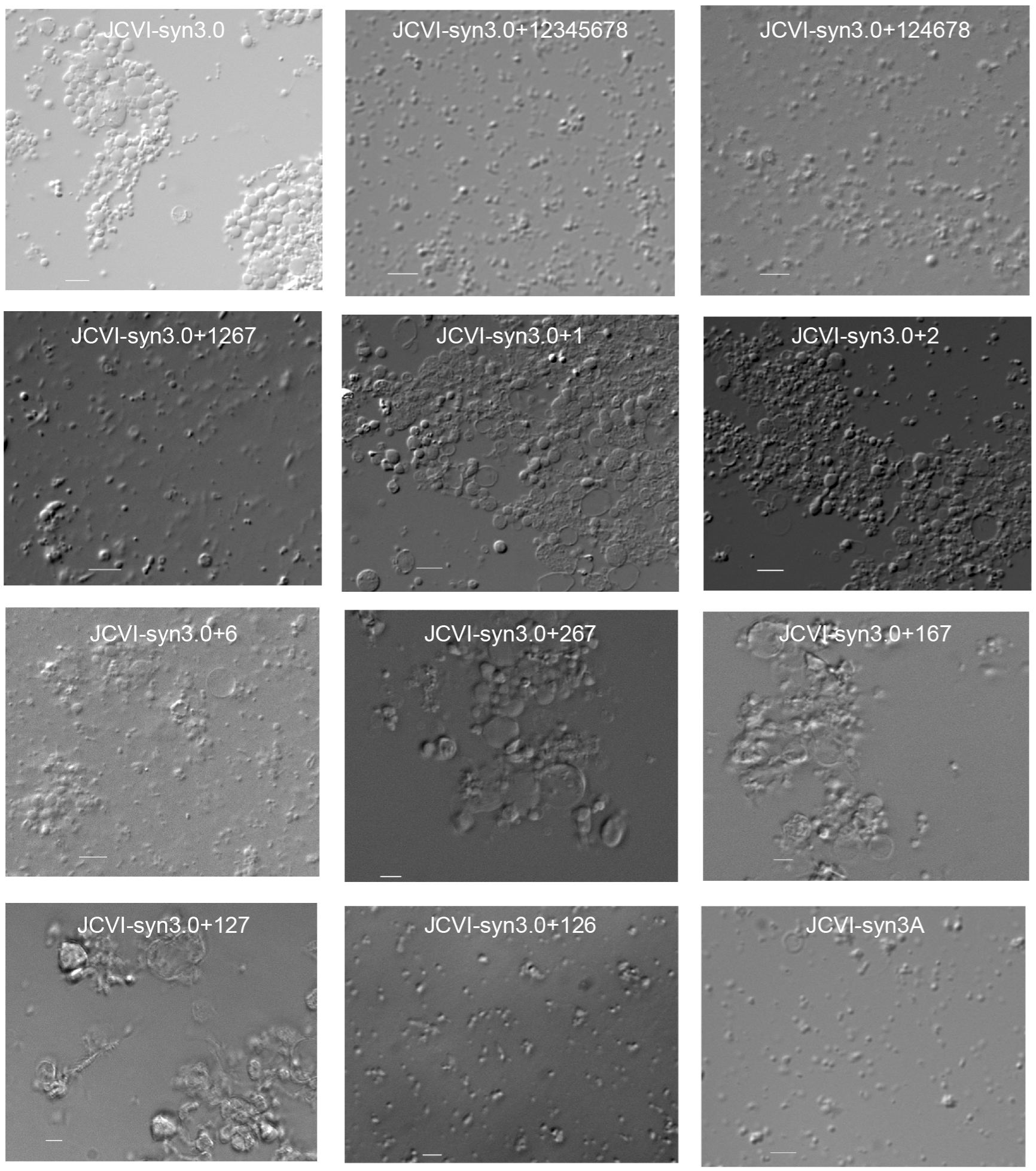
Addition of genes or gene clusters to JCVI-syn3.0. Addition of cluster 1 (*520-522*), cluster 2 (*527*), and cluster 6 (*602,604,605*) was necessary to restore a nearly normal morphology in JCVI-syn3.0+126. These same strains are listed in Fig 5C of the main text. Strains were classified as normal morphology or pleomorphic by scanning samples for large pleomorphic forms and imaging them, if present. Therefore, these images are representative of a much greater number of cells, and the classification as normal morphology is stringent.

**Figure S4B.**
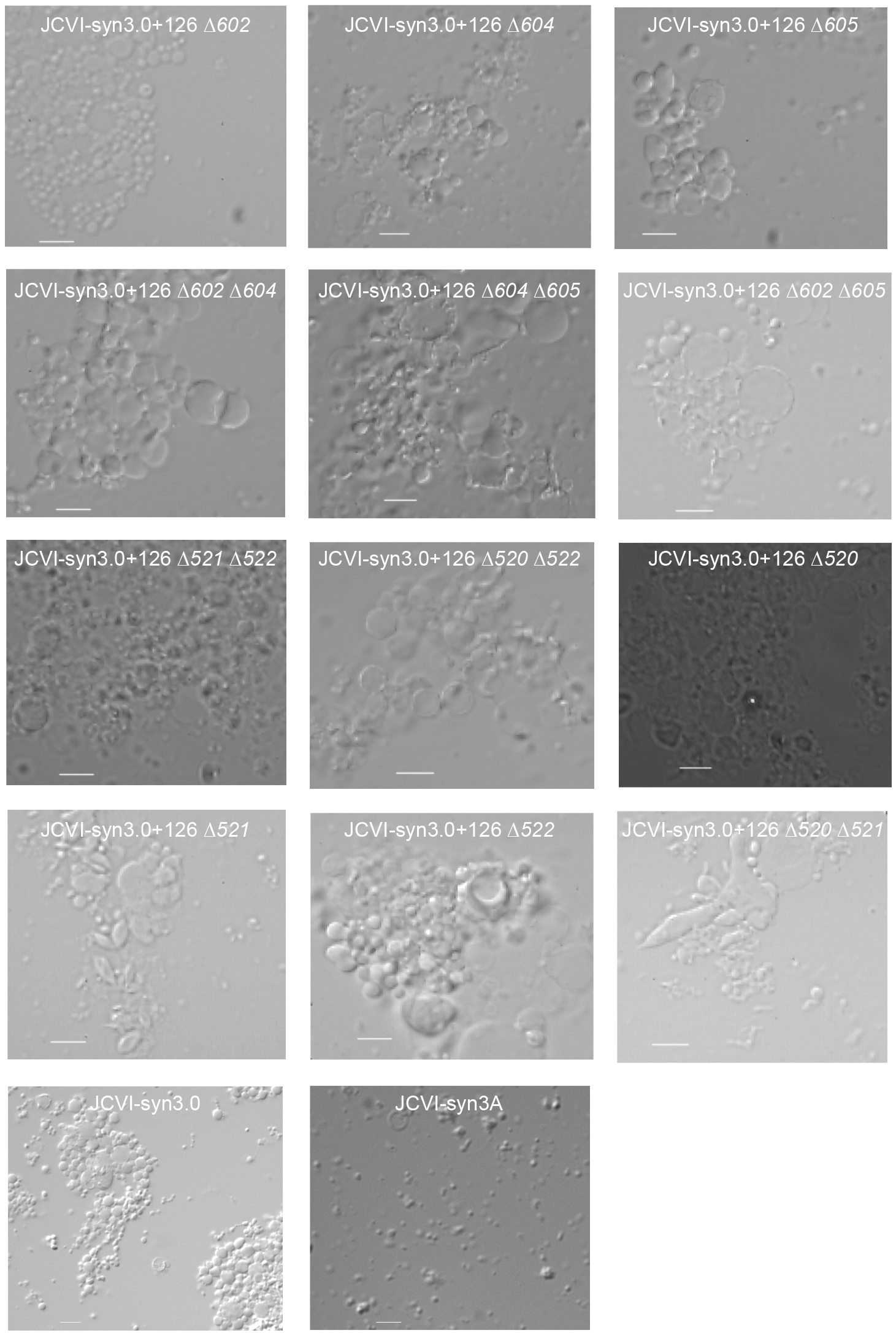
Deletion of genes or gene pairs from JCVI-syn3.0+126. Deletion of any of the seven genes (*520, 521, 522, 527, 602, 604*, or *605*) from JCVI-syn3.0+126 caused loss of the nearly normal morphology.

**Figure S4C.**
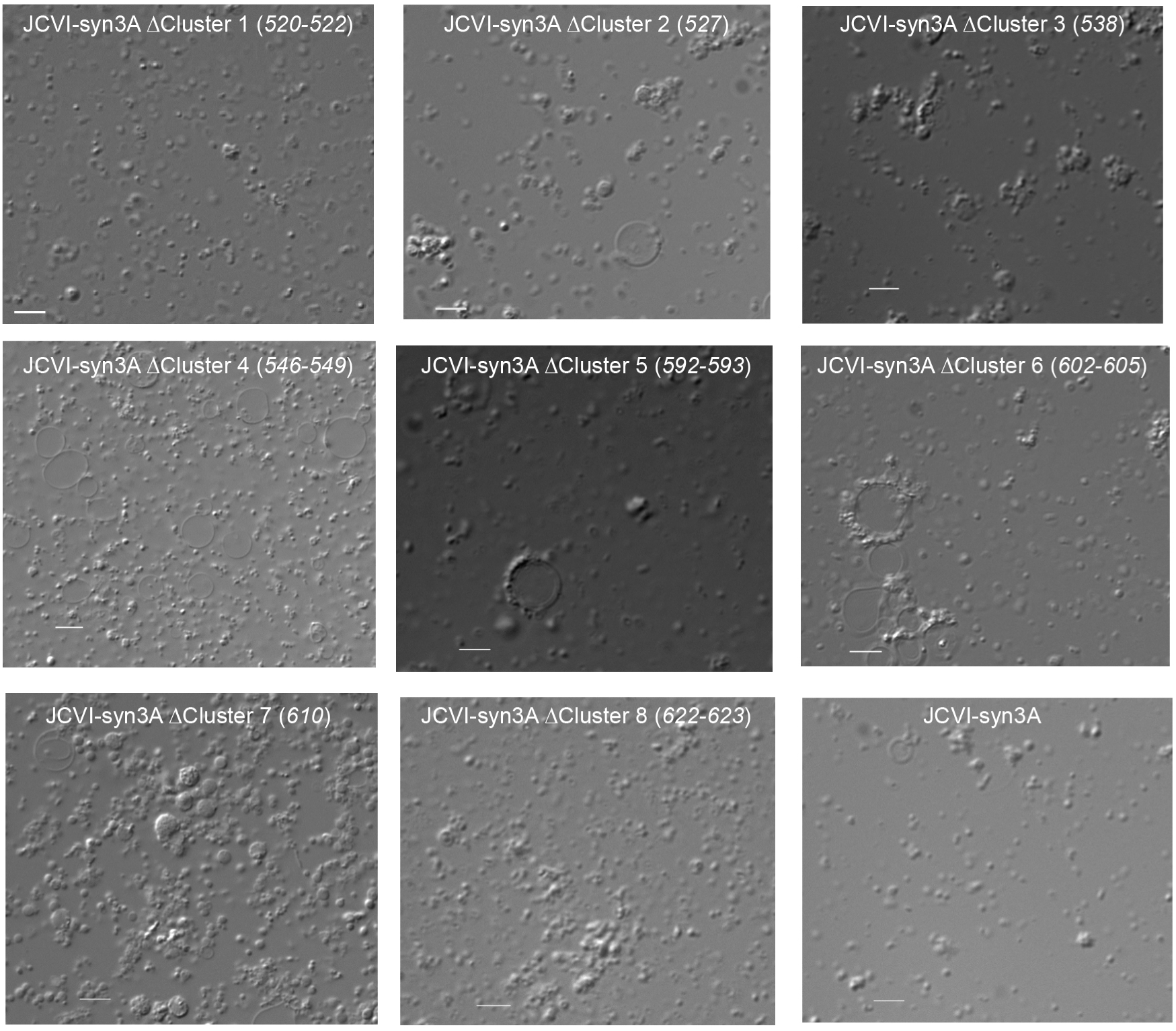
Deletion of genes or gene clusters from JCVI-syn3A. Surprisingly, JCVI-syn3A retained a nearly normal morphology after the deletion of cluster 1 (*520-522*), despite the requirement for cluster 1 for the nearly normal morphology in JCVI-syn3.0+126.

## Notes

### Competing Interest Statement

The authors have declared no competing interest.

